# High-phytate/low-calcium diet is a risk factor for crystal nephropathies, renal phosphate wasting, and bone loss

**DOI:** 10.1101/816512

**Authors:** Ok-Hee Kim, Carmen J. Booth, Han Seok Choi, Jinwook Lee, Jinku Kang, June Hur, Hyung Jin Choi, Hyeonjin Kim, Joong-Hyuck Auh, Jung-Wan Kim, Ji-Young Cha, Young Jae Lee, Cheol Soon Lee, Cheolsoo Choi, Jun-Young Yang, Seung-Soon Im, Dae Ho Lee, Sun Wook Cho, Young-Bum Kim, Kyong Soo Park, Young Joo Park, Byung-Chul Oh

## Abstract

Phosphate overload contributes to mineral bone disorders associated with crystal nephropathies. Phytate, the major form of phosphorus in plant seeds, is known as an indigestible and negligible in humans. However, the mechanism and adverse effects of high-phytate intake on Ca^2+^ and phosphate absorption and homeostasis are unknown. Here we show that excessive intake of phytate with a low-Ca^2+^ diet fed to rats contributed to the development of crystal nephropathies, renal phosphate wasting, and bone loss through tubular dysfunction secondary to dysregulation of intestinal calcium and phosphate absorption. Moreover, Ca^2+^ supplementation alleviated the detrimental effects of excess dietary phytate on bone and kidney through excretion of undigested Ca^2+^-phytate, which prevented a vicious cycle of intestinal phosphate overload and renal phosphate wasting while improving intestinal Ca^2+^ bioavailability. Thus, we demonstrate that phytate is digestible without a high-Ca^2+^ diet and a risk factor for phosphate overloading and developing crystal nephropathies and bone disease.

## Introduction

Phytate *(Myo*-inositol hexaphosphate) is the main phosphorus storage system in plant seeds such as those of cereal grains, nuts, oilseeds, and legumes (Raboy, 2000; Reddy et al., 1982), accounting for up to 80% of total phosphorous. Phytate strongly chelates essential minerals such as Ca^2+^, Mg^2+^, Fe^2+^, and Zn^2+^ to form mineral-phytate salts because of its six phosphate groups (Kim et al., 2010; Oh et al., 2006). These mineral-phytate salts are known to indigestible by monogastric animals and humans that lack the digestive enzyme phytase, which releases phosphate from mineral-phytate salts (Reddy, 2002). The prevailing hypothesis argues that phytate-derived phosphorus is indigestible, not absorbed in the intestine, and is excreted in the feces (Golovan et al., 2001). This widespread belief led to the development of low-phytate crops and commercial phytases to improve the bioavailability of phytate-bound essential minerals including phosphorus and to improve the nutritional quality of plant seeds (Raboy, 2007). Thus, plant seeds or high-fiber vegetable proteins represent a poor or negligible source of bioavailable phosphate for intestinal absorption compared with animal proteins (Moe et al., 2011; Vervloet et al., 2017). Further, they are not considered to adversely affect intestinal phosphate absorption in humans when diets mainly comprise plant-based, cereal-rich diets (Reddy, 2002). Nonetheless, there are insufficient data indicating the amount of phytate phosphorus that is bioavailable for intestinal absorption and whether specific conditions can modulate the bioavailability and digestibility of phytate.

High-phytate diets cause stronger adverse effects on Ca^2+^, iron, and zinc levels among infants, young children, and pregnant and lactating women when cereal-based foods represent a large percentage of the diet (Al Hasan et al., 2016; Chan et al., 2007). Prior studies of dogs demonstrate that increasing the levels of phytate reduces the intestinal absorption of Ca^2+^ and causes nutritional rickets (Harrison and Mellanby, 1939; Mellanby, 1949). Likewise, increasing dietary phytate is associated with secondary hyperparathyroidism in young adults because of the reduced intestinal absorption of dietary Ca^2+^ (Calvo et al., 1988; Pettifor, 1994). Consistent with previous observations (Ford et al., 1972; Goswami et al., 2000), a subsequent case-control study found that nutritional rickets is more prevalent in children who consume diets comprising low dietary Ca^2+^ and high phytate (Aggarwal et al., 2012). These findings suggest a strong association between high-phytate intake and metabolic bone diseases, including nutritional rickets and osteomalacia. However, unlike the well-defined roles of phytate in mineral deficiency (Raboy, 2000; Reddy et al., 1982), whether dietary phytate can modulate intestinal absorption of phosphate is unknown because of a lack of awareness.

Despite its adverse effects on animal and human health, dietary phytate can be beneficial because it inhibits digestive enzymes that hydrolyze lipids, proteins, and starch (Kumar et al., 2010). For example, high-phytate diets inhibit the absorption of glucose and lipids, leading to lower serum levels of glucose, triglycerides, and total cholesterol in humans (Thompson et al., 1987) and rodents (Kim et al., 2010; Jariwalla et al., 1990; Onomi et al., 2004). Moreover, diets based on vegetable protein lead to lower plasma levels of phosphate and fibroblast growth factor 23 (FGF23), a major regulator of phosphate homeostasis (Razzaque, 2009), compared with those based on meat protein (Moe et al., 2011). Indeed, few studies suggest that phytate diets inhibit Ca^2+^ induced kidney stones (Grases et al., 2000). Unfortunately, these studies fail to contribute a broader understanding of the detrimental effects of excess phytate intake on the functions of bone and renal, two major organs that regulate Ca^2+^ and phosphate homeostasis.

Here we investigated the effects of excessive intake of phytate on Ca^2+^ and phosphate homeostasis in a rat model, as well as in a dietary study of humans. Thus, the goals of this study were to determine how a high-phytate diet affects kidney and bone health, establish whether these changes depend on Ca^2+^ or mineral content, and provide a rationale for restricting excessive phytate intake in patients with kidney and metabolic bone diseases.

## Results

### Phytate-Fed rats develop secondary hyperparathyroidism with hypophosphatemia and azotemia

We previously reported that phytate forms insoluble tricalcium- or tetracalcium-phytate salts *in vitro* over a wide pH range (Kim et al., 2010). To evaluate the systemic effects of dietary phytate, rats (4-week-old male and female Sprague Dawley) were AIN-93G diets supplemented with 0% (control), 1%, 3%, or 5% phytate for 12 weeks (**Figure 1A)**. Based on the molecular weights of Ca^2+^ and phytate in these diets, increased phytate supplementation at a constant Ca^2+^ concentration (0.5 %) gradually decreases the molar ratio of Ca^2+^/phytate (**Figure 1B**). Rats fed higher levels of phytate (3% and 5%) exhibited decreased growth rates, as well as reduced fat-free lean mass and whole-body fat despite similar daily food intake compared with those of the control (**Figure 1C–F)**. We reasoned therefore that the decreased growth rates were caused by decreased bioavailability of dietary Ca^2+^ associated with increased phytate supplementation.

**Figure 1.**
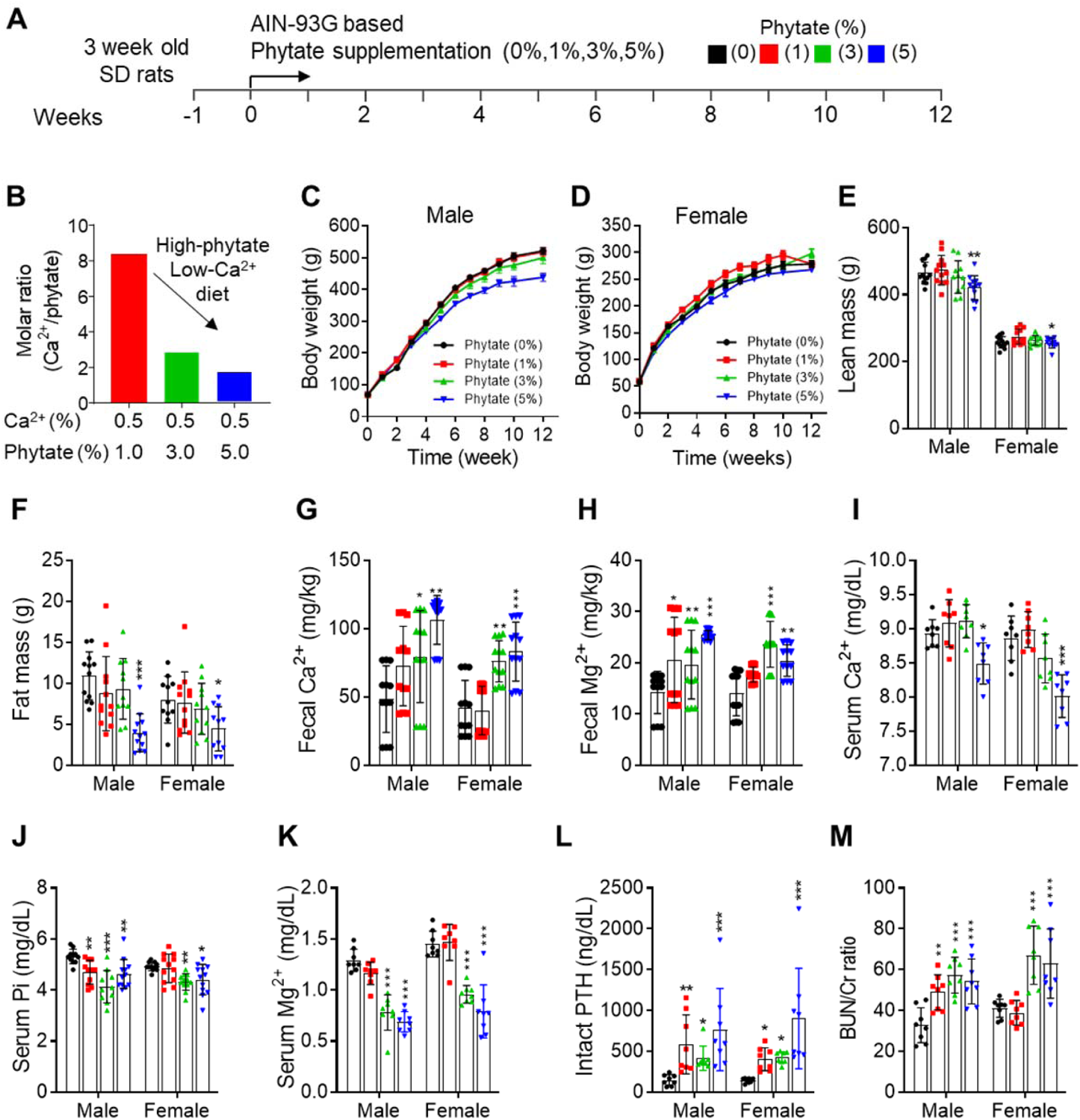
Phytate-Fed rats develop secondary hyperparathyroidism with hypophosphatemia and azotemia. **(A)** Experimental design. Four-week-old male and female Sprague Dawley rats (n=12 per diet) were fed a diet consisting of 0% (control), 1%, 3%, or 5% phytate for 12 weeks. Rats were sacrificed at the end of the experiment. (**B)** Molar ratio of Ca^2+^ to phytate content. (**C, D)** Mean body weights of male (**C**) and female (**D**) rats during the 12-week study. (**E, F)** Mean whole-body lean mass (**E**) and fat mass (**F**) in rats fed AIN-93G diets supplemented with 0%, 1%, 3%, or 5% phytate for 12 weeks. Whole-body lean mass and fat mass were measured using a nuclear magnetic resonance spectrometer. (**G, H)** Fecal excretion of Ca^2+^ (**G**) and Mg^2+^ (**H**) by rats fed a control or phytate-supplemented diet (n=12 per group) for 3 weeks. (**I-M)** Serum levels of Ca^2+^ (**I**), phosphate (**J**), Mg^2+^ (**K**), intact parathyroid hormone (PTH) (**L**), and blood urea nitrogen (BUN) (**M**) in rats fed a control or phytate-supplemented diet for 12 weeks. Serum PTH (intact) or 25(OH)D_3_ levels were measured using an enzyme-linked immunosorbent assay (ELISA) or a radioimmunoassay, respectively. All comparisons were analyzed by one-way ANOVA, *P* values were corrected using Dunnett’s multiple comparisons test. All data are presented as the standard deviation (SD) for each group (n=8–12 per group). \**P* < 0.05, **P < 0.01, ***P < 10^−3^ compared with controls.

To address this possibility, we used inductively coupled plasma mass spectroscopy (ICP-MS) to analyze fecal minerals. We found that dietary phytate led to the inhibition of intestinal absorption of essential minerals such as Ca^2+^, magnesium, and iron in a concentration-dependent manner (**Figure 1G–H–figure supplement 1A**). Serum levels of Ca^2+^ remained within the normal range except in rats fed 5% phytate, associated with slightly reduced serum Ca^2+^ (**Figure 1I**). High-phytate-fed rats developed hypophosphatemia (**Figure 1J**) and hypomagnesemia (**Figure 1k**). Parathyroid hormone (PTH) levels markedly increased (5.4- to 11.7-fold) in the high-phytate groups by week 5 (**figure supplement 1B)** and remained elevated (2.8- to 6.4-fold) to week 12 (**Figure 1L**) in all phytate-fed rats because of sustained inhibition of intestinal Ca^2+^ absorption. These results suggest that PTH plays a role in the development of hypophosphatemia in high-phytate-fed rats.

Serum levels of creatinine (Cr) remained within the normal range except in rats fed 5% phytate, where Cr levels were increased compared with controls (**figure supplement 1C**). In contrast, serum levels of blood urea nitrogen (BUN) showed a marked increase in rats fed high levels of phytate (3% and 5%) compared with controls (**Figure 1M**). The BUN/Cr ratio is a common laboratory value used to diagnose azotemia and acute kidney injury (Uchino et al., 2012). Therefore, we determined the BUN/Cr ratio in phytate-fed rats. Rats exhibited a large increase in the BUN/Cr ratio in all phytate groups (**figure supplement 1D**). These results indicate that a high-phytate diet may lead to the development of azotemia.

### Phytate-fed rats develop crystal nephropathies

To identify the cause of azotemia, we analyzed T2-weighted magnetic resonance images (MRI) of phytate-fed rats. MRI analysis revealed that rats fed 5% phytate had multiple renal cysts at the corticomedullary junction (**Figure 2A–figure supplement 2A**). The kidneys of these rats exhibited pronounced renomegaly with pitted, irregular, and granular cortices and medullar morphology (**figure supplement 2B–C**). The histopathological changes in the kidneys were evaluated while blinded to the experimental group and scored for overall severity of renal injury, presence and severity of renal injury, Ca^2+^ deposits, and fibrosis. The severity of renal injury was greatest in female rats fed 3% or 5% phytate and in male rats fed 5% phytate, whereas controls lacked significant pathological changes.

**Figure 2.**
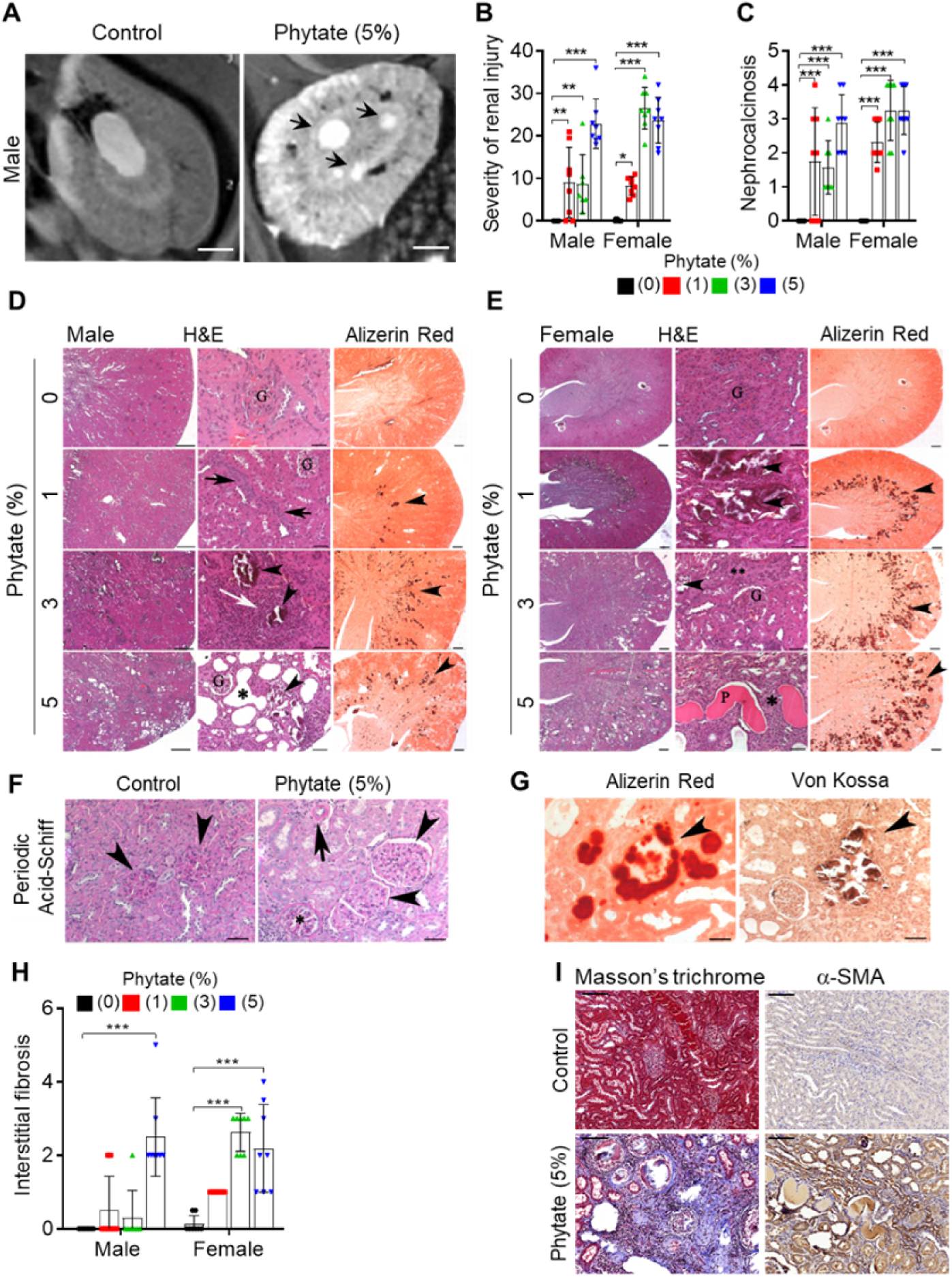
Phytate-fed rats develop crystal nephropathies. **(A)** Representative T2-weighted magnetic resonance images (MRIs) of kidneys from male rats fed a 5% phytate or control diet. The red arrow points to kidney cysts in a phytate-fed rat. Scale bars, 250 µm. (**B–E**) Kidney injury was examined by staining with hematoxylin and eosin, Alizarin Red S (AR), von Kossa (VK), or periodic acid-Schiff (PAS) staining. Scale bars, left 500 µm, center 50 µm, right 500 µm. Granular material within the tubules (nephrocalcinosis) (**D, E**, arrowheads) was present in the kidneys of phytate-fed rats visualized using Alizarin Red S staining (**D, E**, right panels). Pathologic changes included tubular regeneration (**D**, black arrows), inflammation (**D**, white arrow), ectatic tubules (**D**, E,*), protein casts (**E**, P), interstitial fibrosis (E**), and small shrunken glomeruli (**E**, lower panel G). (**F**) Representative PAS-stained sections from control (left) or phytate-fed (right) groups. (**G**) Calcification was detected using Alizarin Red S (left) and VK (right) stains. VK staining for Ca^2+^ was similar to that of Alizarin Red S (arrowheads). In phytate-fed rats with severe pathologic changes, glomeruli were often large (5% phytate, arrowheads) with increased PAS-positive material in the mesangium, Bowman’s capsule of the glomeruli (*), and in the basement membrane of the tubules (arrow). Kidneys from control rats were unremarkable. Scale bar, 50 µm. (**H, I**) Interstitial fibrosis was further detected by staining with Masson’s trichrome (**H**) or immunohistochemistry using antibodies against α-smooth muscle actin (**I**). Scale bar, 100 µm. All comparisons were analyzed by one-way ANOVA, *P* values were corrected using Dunnett’s multiple comparisons test. All data are presented as the SD for each group (n=8 per group). \**P* < 0.05, **P < 0.01, ***P < 10^−3^ compared with controls.

The presence and severity of nephrocalcinosis matched the severity of renal injury scores (**Figure 2B–C**). Pathological changes in phytate-fed rats were consistent with mild to severe crystal nephropathies (Mulay and Anders, 2017; Shavit et al., 2015), including interstitial fibrosis, interstitial inflammation, tubular degeneration (ectasia, nephrocalcinosis, and protein casts), and enlarged or shrunken glomeruli (**Figure 2D–F–figure supplement 2C–F**). Alizarin Red S and von Kossa staining confirmed nephrocalcinosis, showing that the kidneys from phytate-fed rats contained extensive medullary Ca^2+^ deposits (**Figure 2G**). Interstitial fibrosis in phytate-fed rats was further confirmed using Masson’s trichrome staining and immunohistochemical analysis of α-smooth muscle actin, a marker of fibrosis (**Figure 2 H–I**). Together, the histopathologic data demonstrate that a high-phytate diet resulted in marked renal injury with overt nephrocalcinosis and crystal nephropathy.

### Phytate-fed rats develop severe bone loss

Growth retardation and metabolic bone disease are common in children and adolescents with renal disease (Wesseling-Perry et al., 2012). To determine if the phytate-fed rats developed skeletal changes and bone pathology coincident with renal disease, we performed dual-energy X-ray absorptiometry (pDEXA) and histological analyses of excised femora. Phytate-fed rats exhibited marked decreases in the rates of bone accrual with decreased bone mineral density (BMD) in a concentration-dependent manner compared with controls (**Figure 3A**). μ-CT images revealed concomitant increases in the bone marrow cavity space in phytate-fed rats compared with controls (**Figure 3B**). Further, phytate-fed rats exhibited significantly decreased trabecular bone volume (BV/TV), trabecular bone surface (BS/TV), trabecular number (Tb.N), and significantly increased trabecular space (Tb.Sp) compared with controls (**Figure 3C)**. Consistent with the trabecular bone phenotypes, phytate-fed rats had markedly reduced cortical bone area (Tt.Ar) and increased cortical area fractions (Ct.Ar/Tt.Ar) (**Figure 3D**).

**Figure 3.**
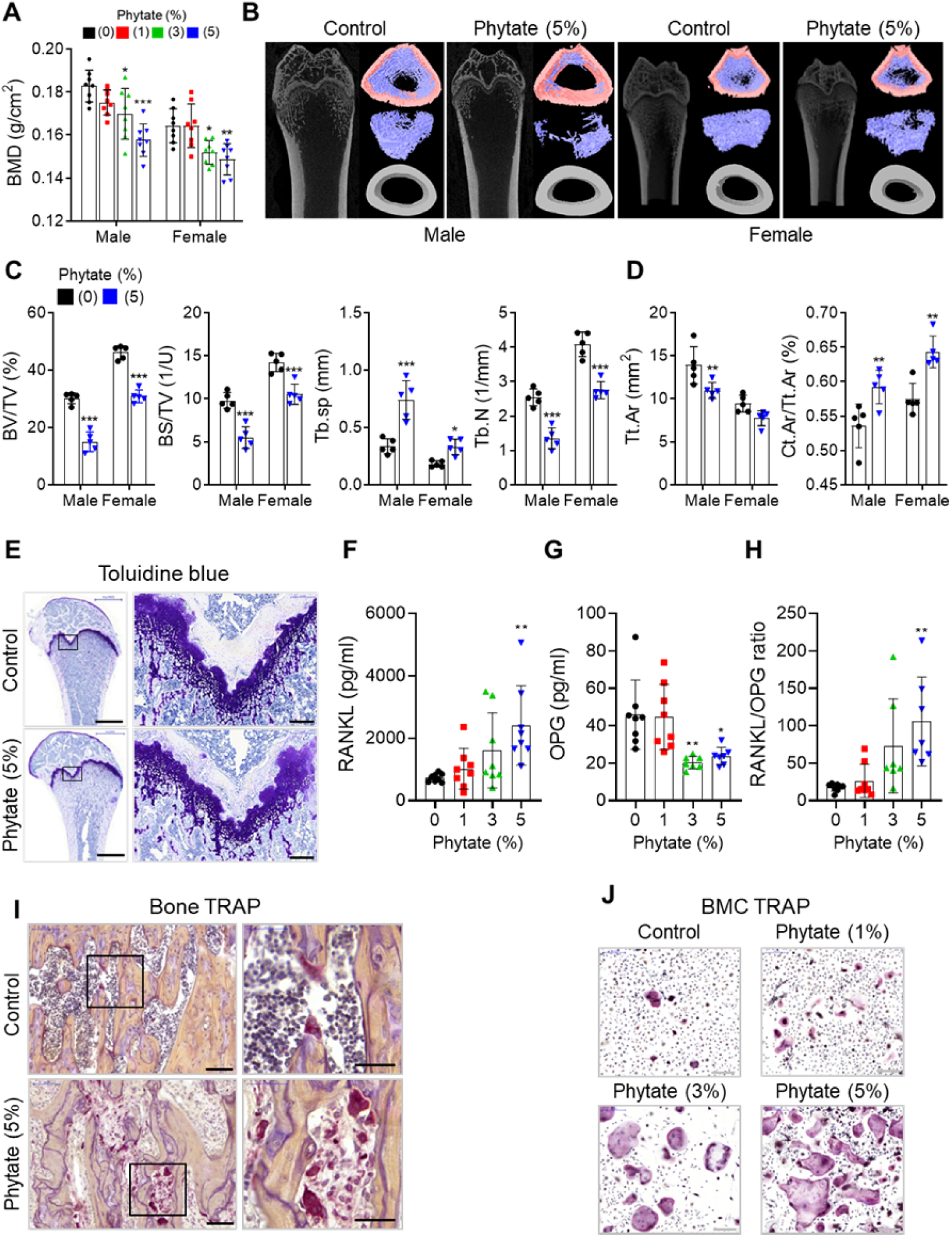
Phytate-fed rats develop severe bone loss. **(A)** Bone mineral density (BMD) was examined using peripheral dual-energy X-ray absorptiometry (pDEXA) of whole femora from rats fed control or phytate-supplemented diets (n=8 per group). (**B)** Three-dimensional reconstructed μ-CT images of distal femora of rats fed control or phytate-supplemented diets. (**C)** Analysis of secondary spongiosa of the distal femur: bone volume fractions (BV/TV), bone surface density (BS/TV), trabecular separation (Tb.Sp), and trabecular number (Tb.N). (**D)** Analysis of the cortical bone of the distal femur: total cross-sectional area inside the periosteal envelope (Tt.Ar) and cortical area fraction (Ct.Ar/Tt.Ar). (**E)** Representative Toluidine blue-stained sections of decalcified femora from female rats fed control or phytate-supplemented diets. Scale bars, left 1000 µm, right 100 µm. (**F–H)** Serum levels of soluble RANKL (**F**), OPG (**G**), and sRANKL/OPG ratio (**H**) in female rats fed control or phytate-supplemented diets. Serum sRANKL and OPG were measured using an ELISA (n=8 per group). (**I)** Representative images of tartrate-resistant acid phosphatase (TRAP)-stained, EDTA-decalcified femur sections of rats fed control (upper) or phytate-supplemented (lower) diets. Red reaction product represents TRAP^+^ multinucleated osteoclasts. Scale bars, left 500 µm, right 100 µm. (**J**) Bone marrow macrophages of rats fed control or phytate-supplemented diets for 12 weeks were cultured with RANK and macrophage colony-stimulating factor for 5 days. Osteoclasts were stained to detect TRAP activity. Scale bars, 200 µm. Comparisons were analyzed by unpaired Student’s *t* test (**C-D**) or one-way ANOVA (**F-H**), *P* values were corrected using Dunnett’s multiple comparisons test. All data are presented as the mean ± SD of each group (**C-D**, 0% and 5% phytate, n = 5 per group). (F-H, n = 8 per group) \**P* < 0.05, **P < 0.01, ***P < 10^−3^ compared with controls.

Male and female rats exhibited the same skeletal alterations, so we characterized bone histopathology of the latter. Bone sections stained with Toluidine blue showed that high-phytate diets decreased cartilage growth-plate thickness and bone osteoids (**Figure 3E**), consistent with skeletal alterations. Serum levels of receptor activator of nuclear factor kappa-B ligand (RANKL), a key component of osteoclastogenesis (Kong et al., 1999), were markedly increased in high-phytate-fed rats compared with controls (**Figure 3F**). The levels of osteoprotegerin (OPG), a RANKL decoy receptor that inhibits osteoclast differentiation and activation (Lacey et al., 1998), were significantly reduced (**Figure 3G**), and the sRANKL/OPG ratio was significantly increased in phytate-fed rats (**Figure 3H**). These data suggest a systemic acceleration of osteoclastogenesis in response to dietary phytate. Analysis of bone sections stained for tartrate-resistant acid phosphatase (TRAP) revealed that consumption of the high-phytate diet resulted in increased TRAP^+^ multinucleated cells (**Figure 3I**). Further, bone marrow-derived stromal cell cultures exhibited a significant concentration-dependent increase in the number of giant TRAP^+^ multinucleated osteoclasts (**Figure 3J**). Together, the data led us to posit that a high-phytate diet contributed to severe bone loss.

### High-phytate diets dysregulate mineral and phosphate metabolism in humans

To determine if our findings using rats were applicable to humans, we conducted a non-comparative pilot study to evaluate the short-term effects of phytate on six healthy premenopausal women (aged 23– 34 years, body mass index, 17.7–25.8 kg/m^2^) (**Figure 4A**). The 12-day trial consisted of three consecutive diet cycles (four days in each cycle) of (1) polished white rice (WR, 0.35% phytate); (2) brown rice (BR, 1.07% phytate); and (3) brown rice with rice bran (BR+RB, 2.18% phytate). At the end of each diet cycle, 16-h fasting blood and 24-h urine samples were collected from each participant. Serum levels of Ca^2+^, magnesium, and phosphate remained within normal range in all three diets, although high dietary phytate intake was associated with slightly decreased serum phosphate levels (**Figure 4B–C**). Results of other biochemical parameters are summarized in **table supplement 1**.

**Figure 4.**
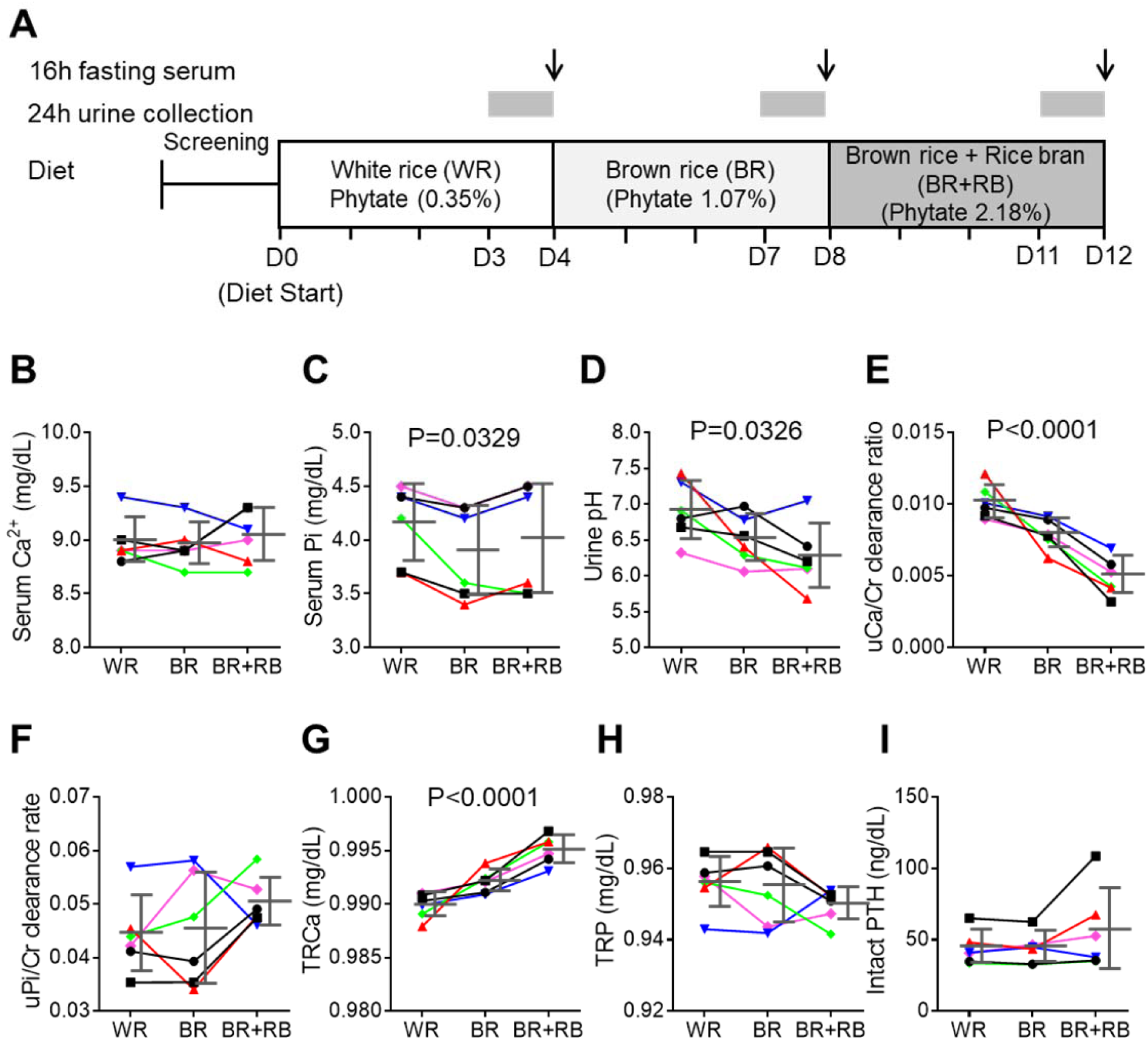
High-phytate diets dysregulate mineral and phosphate metabolism in humans. **(A)** Experimental design (6 healthy female volunteers). (**B**,**C)** Time course of serum levels of Ca^2+^ (**B**) and phosphate (**C**) for three different phytate doses. (**D–H)** Kinetics of urine pH (**D**), 24-h urine Ca^2+^/Cr ratio (**E**), 24-h urine Pi/Cr ratio (**F**), renal tubular Ca^2+^ reabsorption (TRCa, **G**), and renal tubular phosphate reabsorption (TRP, **H**) for three different phytate doses. (**I**) Time-course analysis of serum levels of intact PTH for different phytate doses. Data are presented as the mean ± SEM for each dose (n 6 per dose). Statistical significance were tested by repeated measure ANOVA between subjects and groups. Data are presented as the mean ± SD for each dose (n 6 per dose).

Urinary pH levels decreased in a dose-dependent manner as the intake of phytate increased (**Figure 4D**). Higher dietary phytate intake resulted in a marked decrease in the 24-h urinary Ca^2+^ creatinine clearance rate (**Figure 4E**), while the 24-h urinary phosphate creatinine clearance rate was only slightly increased (**Figure 4F**). These findings are consistent with the results of the tubular reabsorption rate, where high dietary phytate increases the renal tubular Ca^2+^ reabsorption rate (**Figure 4G**) but decreases the renal tubular phosphate reabsorption rate (**Figure 4H**). These metabolic characteristics are likely secondary to the increased levels of PTH in response to high levels of dietary phytate intake (**Figure 4I**). Further, high dietary phytate decreased the serum levels of FGF23 (**figure supplement 3A**). Thus, we propose that the high dietary phytate intake dysregulates mineral and phosphate metabolism in humans in a manner consistent with the findings of our rat studies.

### Ca^2±^ supplementation protects phytate-fed rats from renal phosphate wasting via indigestible Ca^2+^-phytate salts

Our study showed that diets containing >2%–3 % phytate dysregulate Ca^2+^ and phosphate homeostasis in rats as well as in humans. Thus, when we calculated the daily amount of consumption of 3% phytate in rats, we found that administering 150–300 mg/kg phytate per day was harmful to rats. This daily dose range of phytate may correspond to 24.2–48.4 mg/kg body weight of humans (1,451–2,900 mg/60 kg) according to a simple practice guide for dose conversion between animals and humans (Nair and Jacob, 2016). This calculated daily dose range of dietary phytate may account for the >4-fold higher average intake compared with those in the United State and United Kingdom (631–746 mg) (Ellis et al., 1987).

Phytate binds Ca^2+^ with high affinity at physiological pH (Kim et al., 2010; Reddy, 2002). Consequently, the detrimental effects of phytate only occur when a diet poor in trace elements or Ca^2+^ is consumed (Raboy, 2007; Humer et al., 2015). We asked therefore whether increasing the molar ratio of Ca^2+^/phytate can modulate the bioavailability and digestibility of phytate *in vivo*. To address this question, we fed female rats a diet of 3% phytate supplemented with high (1%) Ca^2+^ (high phytate with high Ca^2+^ [HP-HCa^2+^]) to increase the molar ratio of Ca^2+^/phytate and Ca^2+^m bioavailability compared with 3% phytate supplemented with low (0.5%) Ca^2+^ (high phytate with low Ca^2+^ [HP-LCa^2+^]) (**Figure 5A**). The growth rates of rats fed the HP-HCa^2+^ diet were decreased and not associated with significant changes in serum levels of Ca^2+^ despite slightly higher food and water intake compared with controls (**figure supplement 4A–D**). Moreover, rats fed the HP-HCa^2+^ diet completely normalized their serum levels of phosphate and BUN, as well as the BUN/Cr ratio compared with rats fed the HP-LCa^2+^ diet (**Figure 5B–C–figure supplement 4E**), indicating that supplementation with high Ca^2+^ ameliorated hypophosphatemia and azotemia.

**Figure 5.**
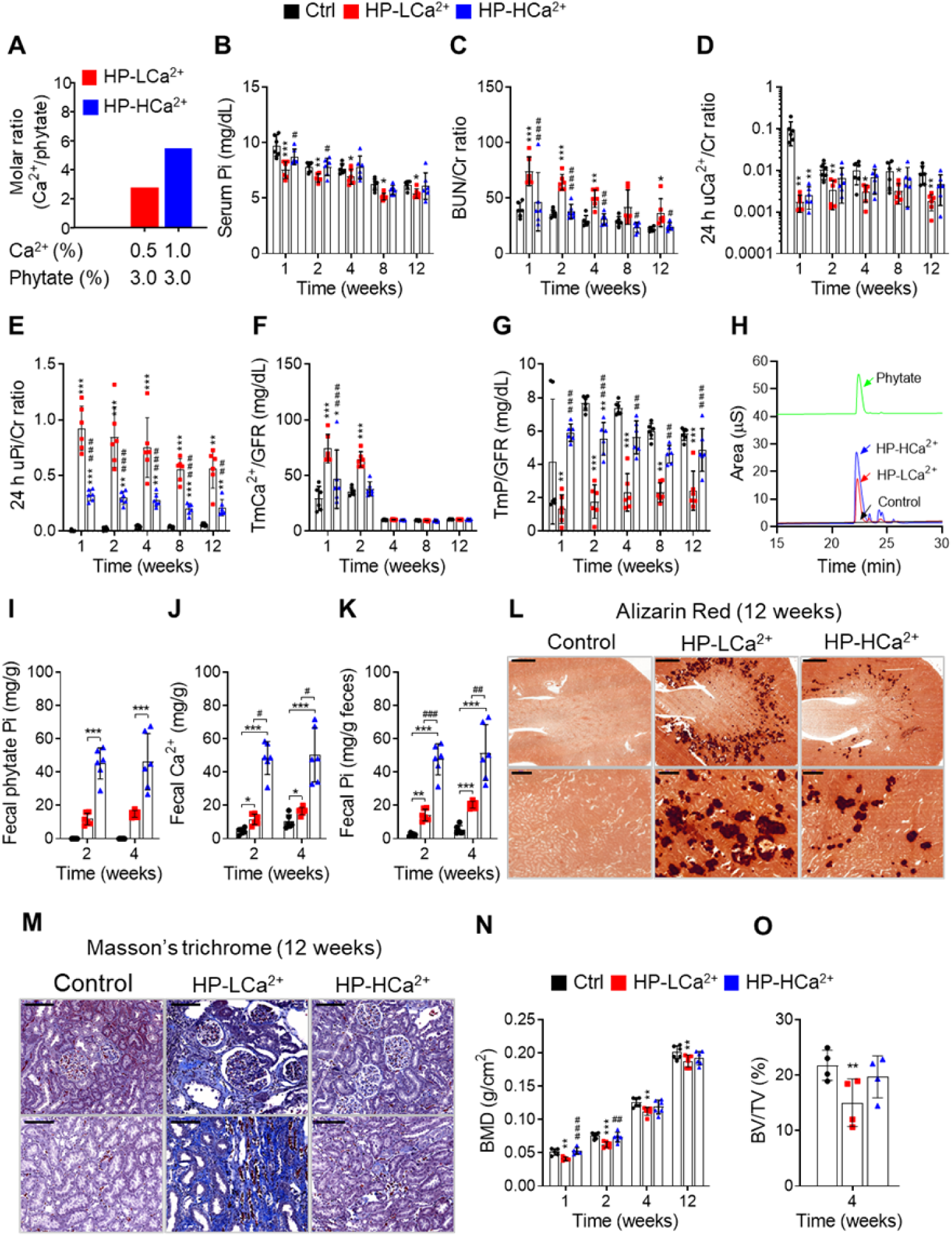
Ca^2+^ supplementation protects phytate-fed rats from renal phosphate wasting disorders, crystal nephropathies, and bone loss via indigestible Ca^2+^-phytate salts. **(A)** Molar ratio of Ca^2+^ to phytate content when the diet was supplemented with 1.0% Ca^2+^. (**B–C)**, Time-course analysis of serum levels of Pi (**B**) and BUN/Cr ratio (**C**) in rats fed control, HP-LCa^2+^, and HP-HCa^2+^ diets for 12 weeks. (**D–G)** Time-course analysis of 24-h urine Ca^2+^/Cr ratio (**D**), 24-h urine Pi/Cr ratio (**E**), ratio of tubular maximum reabsorption rate of renal tubular Ca^2+^ to glomerular filtration rate (**F)** TmCa^2+^/GFR, and ratio of tubular maximum reabsorption of renal tubular phosphate to glomerular filtration rate (**G)** TmP/GFR in in rats fed control, HP-LCa^2+^, and HP-HCa^2+^ diets for 12 weeks. (**H–K)**, Fecal phytate was quantified using HPIC (**H, I**), and fecal Ca^2+^ and Pi were analyzed using ICP-MS (**K**). Time-course analysis of fecal phytate (**I**), Ca^2+^ (**J**), and phytate Pi (**K**) in rats fed control, HP-LCa^2+^, and HP-HCa^2+^ diets. Fecal phytate phosphorus was calculated as follows (ICP-MS analysis:[Phytate Pi] = (([Pi] from rats fed a HP-LCa^2+^ or HP-HCa^2+^ diets) - ([Pi] from rats fed control diet)). (**L, M**) Representative Alizarin Red S staining (**L**) and Masson’s trichrome staining (**M**) of kidney sections of rats fed control, HP-LCa^2+^, and HP-HCa^2+^ diets for 12 weeks.). (**N, O)** Kinetics of BMD (**N**) and bone volume fractions (**O**, BV/TV) from rats fed control, HP-LCa^2+^, and HP-HCa^2+^ diets. All comparisons were conducted using two-way ANOVA with Tukey’s post hoc multiple comparison testing. All data are presented as the mean ± SD of each group (n=4-6 per group). *P < 0.05, **P < 0.01, ***P < 10^−3^, compared with controls. ^#^P<0.05, ##P<0.01, ^###^P<10^−3^, compared with a HP-LCa^2+^ diet groups.

To identify the mechanism of phytate-induced hypophosphatemia, we collected 24-h urine samples from phytate-fed rats. Rats fed the HP-LCa^2+^ diet had markedly decreased urine pH (**figure supplement 4F**) without significant differences in urine volume (**figure supplement 4G**), marked decreases in the 24-h urinary Ca^2+^/Cr ratio, and a marked increase in the 24-h urinary Pi/Cr ratio compared with controls (**Figure 5D–E**). Further, rats fed the HP-LCa^2+^ diet had markedly higher renal tubular reabsorption of Ca^2+^ (TmCa^2+^/GFR), with lower renal tubular reabsorption of phosphate (TmP/GFR) compared with controls (**Figure 5F–G**). We surmised that the high-phytate diet decreased urinary Ca^2+^ excretion while increasing renal phosphate wasting. Impaired renal Ca^2+^ and phosphate reabsorption are restored in rats fed the HP-HCa^2+^ diet (**Figure 5F–G**).

High-performance ion chromatographic (HPIC) analysis of feces demonstrated that rats fed the HP-HCa^2+^ diet excreted an approximately 3-fold higher level of fecal phytate compared with rats fed the HP-LCa^2+^ diet (**Figure 5H–I**). ICP-MS analysis, performed after acid hydrolysis of fecal phytate to generate phosphate, further confirmed that rats fed the HP-HCa^2+^ diet exhibited an approximately 3.8-fold greater increase in Ca^2+^ and phosphate excretion compared with rats fed the HP-LCa^2+^ diet (**Figure 5J–K**), confirming findings of increased fecal excretion of undigested Ca^2+^-phytate in rats fed the HP-HCa^2+^ diet. Thus, we conclude that rats fed the HP-HCa^2+^ diet accumulated unabsorbed and indigestible Ca^2+^-phytate salts in the intestine while decreasing the intestinal phosphate available for absorption, indicating that the endogenous enzyme phytase was unable to hydrolyze Ca^2+^-phytate salts.

Conversely, rats fed the HP-LCa^2+^ diet hydrolyzed and absorbed >68%–74% of the phosphate in the intestine **(Figure 5H–K)** with a concomitant increase in urinary phosphate excretion, thus developing renal phosphate wasting (**Figure 5E–G**). These data indicate that rats that were fed the HP-LCa^2+^ diet exhibited increased intestinal phosphate overloading caused by excessive phytate hydrolysis, which is consistent with evidence indicating that renal excretion of phosphate increases when phosphate intake is excessive (Vervloet et al., 2017). Thus, our findings demonstrate that phytate was digestible and absorbed in the intestine, considering that phytate is not digested and absorbed *in vivo* when it forms Ca^2+^-phytate salts in the presence of high Ca^2+^ concentrations. Further, these findings are consistent with those of our previous *in vitro* studies (Kim et al., 2010).

### Ca^2+^ supplementation protects phytate-fed rats against crystal nephropathies and bone loss

We next examined the morphological and histopathological changes in bones and kidneys when rats were fed the HP-LCa^2+^ diet. Ca^2+^ supplementation protected these rats against the development of the expected severe gross and microscopic changes of renomegaly (**figure supplement 5A–C**) and medullary nephrocalcinosis (**Figure 4l–figure supplement 5D–G**), respectively, revealed by Alizarin Red S staining. Further, rats fed the HP-HCa^2+^ diet expressed lower levels of four key markers (Kronenberg, 2009) of renal damage and fibrosis (*NGAL, COL1A1, COL6A2*, and *MPG*) (**figure supplement 6A**) and exhibited decreased renal fibrosis revealed by Masson’s trichrome staining to detect collagen (**Figure 4M–figure supplement 6B–C**). These data suggest that the HP-LCa^2+^ diet was a key contributor to renal damage and fibrosis. Further, pDEXA and μ-CT analyses of excised femora confirmed that rats fed the HP-HCa^2+^ diet were protected against the expected bone loss and demineralization observed compared with rats fed the HP-LCa^2+^ diet (**Figure 4N–O–figure supplement 6D–G**). Together, these findings support our hypothesis that high dietary phytate intake leads to damage of renal tissues as well as impaired bone mass accrual, while Ca^2+^ supplementation of the high-phytate diet has protective effects.

### Phytate-mediated Ca^2+^ deficiency promotes vitamin D insufficiency and renal phosphate wasting independent of FGF23 expression

To identify the physiological mechanisms of high-phytate diets on vitamin D metabolism, we performed a time-course analysis of the serum levels of PTH, 25(OH)D_3_, and activated D_3_ (1,25[OH]_2_D_3_). First, we found that the serum levels of PTH were consistently elevated in rats fed the HP-LCa^2+^ diet compared with control rats and rats fed the HP-HCa^2+^ diet (**Figure 6A**). Second, we found that in conjunction with high PTH, rats fed the HP-LCa^2+^ diet had decreased serum levels of 25(OH)D_3_ compared with controls (**Figure 6B**), leading to increased serum levels of 1,25(OH)_2_D_3_ (**Figure 6C**). In rats fed the HP-HCa^2+^ diet, the serum levels of 25(OH)D_3_ and 1,25(OH)_2_D_3_ normalized, indicating that Ca^2+^ supplementation inhibited the metabolism of serum 25(OH)D_3_ to 1,25(OH)_2_D_3_ (**Figure 6B–C**). Consistent with these findings, Ca^2+^ supplementation normalized the expression levels of *CYP27B1, CYP24A1*, and vitamin D receptor (*VDR*) (**Figure 6D–F**). Further, our findings, which are consistent with those of a previous study (Clements et al., 1987), demonstrate that phytate-mediated inhibition of intestinal Ca^2+^ absorption as well as increased phosphate overload promoted metabolize of 25(OH)D_3_ to 1,25(OH)_2_D_3_, suggesting that high-phytate diets deplete serum 25(OH)D_3_, which mitigates the harmful effects of phytate.

**Figure 6.**
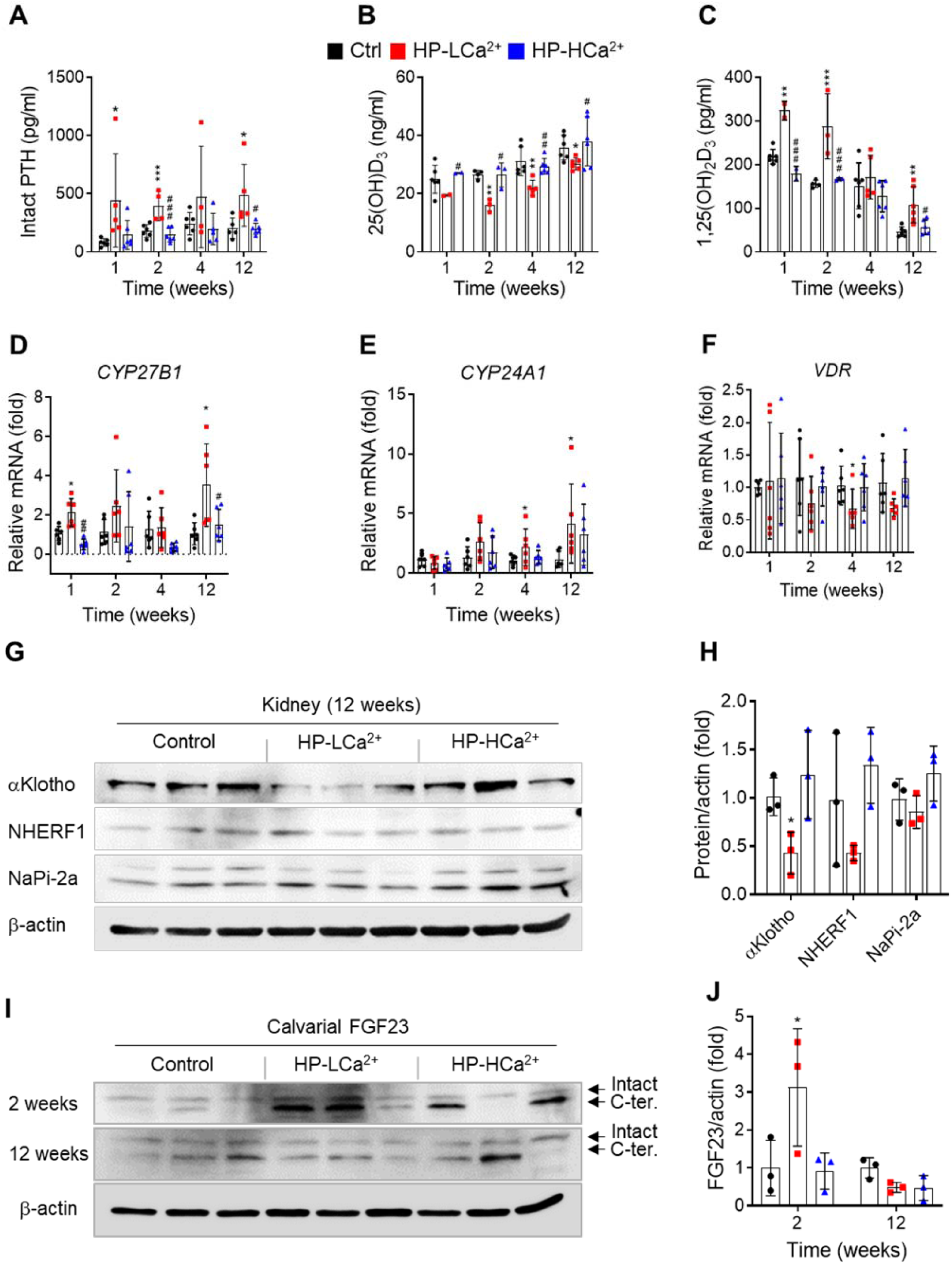
Phytate-mediated Ca^2+^ deficiency promotes vitamin D insufficiency and renal phosphate wasting independent of FGF23 expression. **(A–C)** Time-course analysis of serum levels of intact PTH (**A**), 25(OH)D_3_ (**B**), and 1,25(OH)_2_D_3_ (**C**) in rats fed control, HP-LCa^2+^, and HP-HCa^2+^ diets. For the early measurement of 25(OH)D_3_ (**B**) and 1,25(OH)_2_D_3_, we pooled the sera from 2–3 rats. (**D–F**) Time-course analysis of renal CYP27B1 (**D**), CYP24A1 (**E**), and VDR (**F**) in rats fed control, HP-LCa^2+^, and HP-HCa^2+^ diets. (**G**) Immunoblot analysis of renal αKlotho, NHERF1, NaPi-2a in rats fed control, HP-LCa^2+^, and HP-HCa^2+^ diets. (**H**) The levels of renal proteins were quantified using ImageJ software (NIH, Bethesda, MD, USA). (**I**) Immunoblot analysis of calvarial FGF23 in rats fed control, HP-LCa^2+^, and HP-HCa^2+^ diets. (**J**) The levels of total calvarial FGF23 were quantified using ImageJ software. All comparisons were conducted using two-way ANOVA with Tukey’s post hoc multiple comparison testing. All data are presented as the mean ± SD of each group (n=4-6 per group). *P < 0.05, **P < 0.01, ***P < 10^−3^, compared with controls.

To identify proteins that contributed to phytate-mediated impairment of renal tubular Ca^2+^ and phosphate reabsorption, we analyzed the mRNA levels of key renal genes associated with Ca^2+^ and phosphate homeostasis, including those associated with Ca^2+^, phosphate channels, and regulatory proteins (Mulay and Anders, 2017; Huang et al., 2013). We found that the renal expression of key genes associated with renal phosphate wasting, including Ca^2+^ sensing receptor (CaSR), inositol 1,4,5-trisphosphate receptor 1 (ITPR1), αKlotho, sodium/phosphate cotransporters (NaPi-2a and NaPi-2c), sodium-hydrogen antiporter 3 regulator 1 (NHERF1), and phosphate-regulating neutral endopeptidase X-linked was markedly decreased in phytate-fed rats (**figure supplement 7A**). Each of these genes is associated with the development of nephrocalcinosis and renal phosphate wasting (Shavit et al., 2015; Huang et al., 2013; Karim et al., 2008; Monico and Milliner, 2011).

Immunoblot analysis further confirmed the renal expression of αKlotho was significantly decreased in rats fed the HP-LCa^2+^ diet compared with controls (**Figure 5G–H–figure supplement 7B–C**), suggesting that low levels of renal αKlotho may represent a complementary mechanism that increases serum phosphate levels in response to severe hypophosphatemia in rats fed the HP-LCa^2+^ diet. High serum FGF23 levels in conjunction with elevated levels of PTH contribute to the pathogenesis of renal phosphate wasting and hypophosphatemic rickets (Vervloet et al., 2017; Rodriguez-Ortiz et al., 2012; Walton and Bijvoet, 1975). Therefore, we analyzed the serum levels of FGF23. Unexpectedly, quantitative analysis did not detect significant changes in the serum levels of full-length FGF23 or its C-terminal FGF23 in rats fed the HP-LCa^2+^ diet compared with controls (**figure supplement 7D–E**). Immunoblot and immunohistochemical analyses of sections of the femur revealed that rats fed the HP-LCa^2+^ diet showed decreased the levels of FGF23 in the calvarial bone (**Figure 5I–J–figure supplement 7F**), suggesting the phytate-mediated renal phosphate wasting was mainly caused by high PTH levels independent of FGF23.

## Discussion

Here we show that the HP-LCa^2+^ diet was a significant risk factor for phosphate overloading and renal phosphate wasting associated with increased risks of nephrocalcinosis, renal fibrosis, crystal nephropathy, hypophosphatemia, and bone loss. Our findings support our hypothesis that an HP-LCa^2+^ diet has detrimental metabolic effects on Ca^2+^ and phosphate homeostasis in rats through inhibiting intestinal Ca^2+^ absorption while increasing intestinal phosphate overloading. These events lead to high levels of PTH and low levels of FGF23 and αKlotho. In contrast, supplementation with high Ca^2+^ alleviated phytate-mediated pathogenic defects such as biochemical abnormalities, high PTH levels, nephrocalcinosis, renal fibrosis, vitamin D insufficiency, hypophosphatemia, and bone loss. We identified the dietary conditions that influence the bioavailability of phytate when an excess of Ca^2+^ is added to high-phytate diets. Thus, phytate forms insoluble Ca^2+^-phytate salts in the intestine, where it becomes indigestible and unabsorbable, and ultimately was excreted as undigested Ca^2+^-phytate salts in the feces. Thus, Ca^2+^ supplementation normalized impaired renal tubular Ca^2+^ and phosphate reabsorption by preventing intestinal phosphate overloading caused by phytate hydrolysis and improving intestinal Ca^2+^ bioavailability (**Figure 7**).

**Figure 7.**
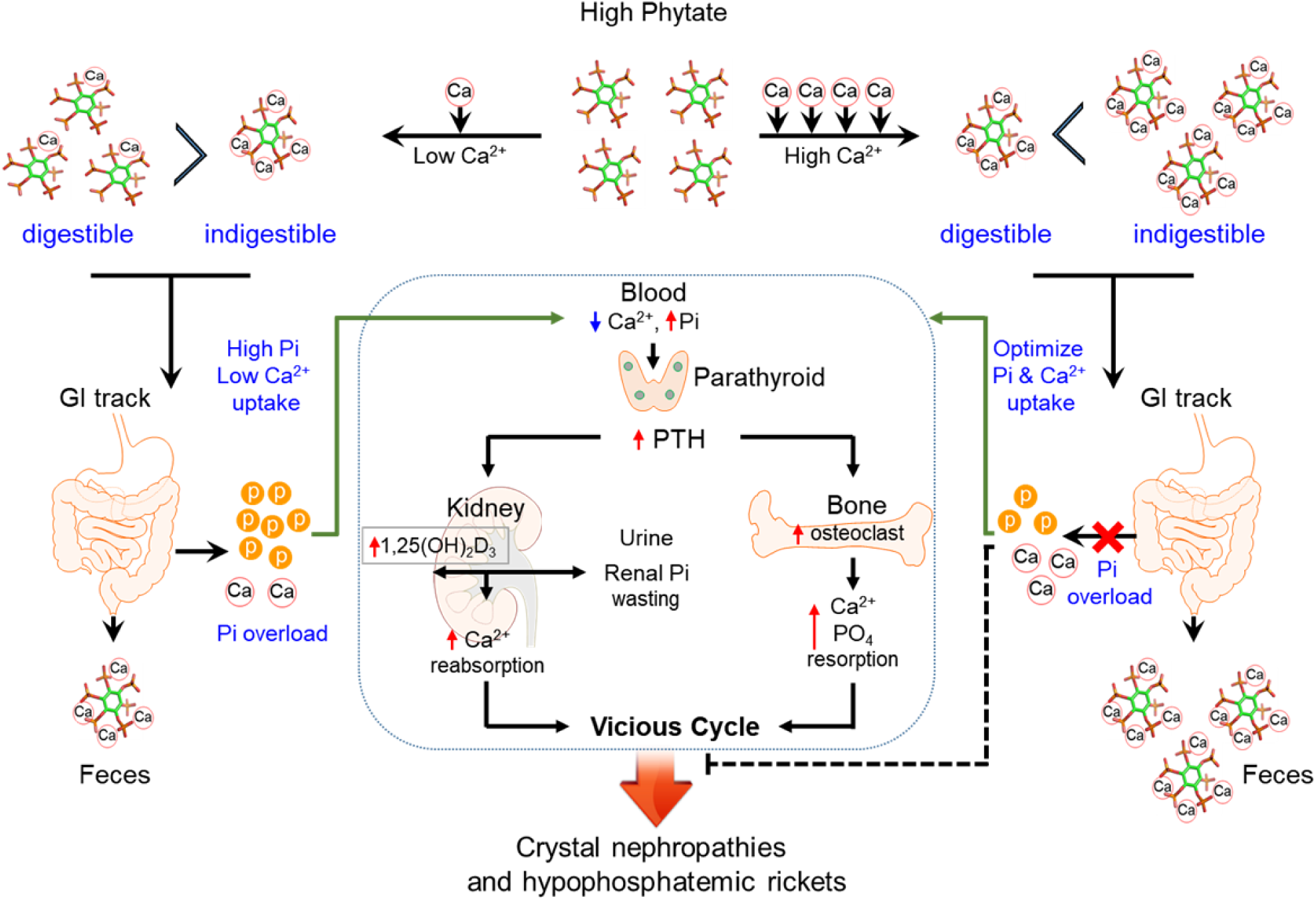
Model for the effects of phytate diets supplemented with low-Ca^2+^ or high-Ca^2+^ on Ca^2+^ and phosphate homeostasis. In the presence of a low-Ca^2+^ diet (Ca^2+^/phytate <5), phytate can adversely affect Ca^2+^ and phosphate homeostasis in normal healthy rats by inhibiting intestinal Ca^2+^ absorption while increasing phosphate overload through phytate hydrolysis in the intestine. These effects lead to crystal nephropathies, severe renal phosphate wasting, and pathologic bone loss via increased PTH and 1,25(OH)_2_D_3_. In contrast, in the presence of a high-Ca^2+^ diet, phytate forms multiple Ca^2+^-phytate salts, which makes them indigestible and unabsorbable in vivo, leading to their excretion in the feces as undigested Ca^2+^-phytate salts. This process prevents a vicious cycle of intestinal phosphate overloading and renal phosphate wasting while improving intestinal Ca^2+^ bioavailability, which eventually alleviates the detrimental effects of excess phytate hydrolysis.

We identified previously unrecognized mechanisms of bioavailability and digestibility of phytate *in vivo*. Thus, HPIC and ICP-MS analyses of feces revealed that phytate is present digestible or indigestible forms *in vivo* depending on the dietary Ca^2+^ content, revealing that phytate is only indigestible and unabsorbable when it forms multiple Ca^2+^-phytate salts. This finding is consistent with data of our *in vitro* studies (Kim et al., 2010), which show that phytate possesses three or four Ca^2+^-binding sites at physiological pH and forms insoluble multiple Ca^2+^-phytate salts. In contrast, when the Ca^2+^/phytate molar ratio is <5 (HP-LCa^2+^ diet), phytate becomes digestible, which leads to phosphate overloading in the intestine leading to renal phosphate wasting and metabolic bone disease. These results may explain why the adverse effects of high-phytate on nutrients are exacerbated when Ca^2+^ intake or mineral content is low (Raboy, 2007; Humer et al., 2015).

Further, the potential toxicity of excess phosphate in the context of mineral bone disorders (MBD) associated with chronic kidney disease (CKD) (Vervloet et al., 2017; Ketteler et al., 2017) and Ca^2+^-phosphate crystals (Kuro-o, 2013), together with our present findings, indicate that a HP-LCa^2+^ diet can be particularly harmful to patients suffering from these conditions. Moreover, dietary phosphate overload can cause tubular damage, hypophosphatemia, Ca^2+^-phosphate crystals; and phosphate overloading never causes persistent hyperphosphatemia unless renal function is critically impaired (Haut et al., 1980). Thus, our study emphasizes that a detailed understanding of the dietary effects and properties of phytate may provide a new opportunity to reduce intestinal phosphate overload and benefit patients with CKD, mineral bone disease, or crystal nephropathies.

Several lines of evidence support our findings that phytate is digestible in the absence of Ca^2+^ or divalent minerals. First, studies of rats fed ^14^C-labeled phytate show that approximately 94% of phytate is absorbed when Ca^2+^ intake is low, whereas 54% of phytate is excreted in the feces when Ca^2+^ intake is high (Nahapetian and Young, 1980), suggesting that phytate absorption is modulated or inhibited by the Ca^2+^ concentration. Second, another study of rats fed ^3^H-inositol phytate shows that 78% of phytate is absorbed and 14% is excreted in the feces. Further, most of the radioactivity represents *myo*-inositol or *myo*-inositol monophosphate in the plasma or urine (Sakamoto et al., 1993), suggesting the phytate is digestible *in vivo*. The biased belief that phytate is indigestible led the authors of the above two studies to conclude that intact phytate is absorbed through the upper intestine, hydrolyzed within mucosal cells before entry into the blood and then released as *myo*-inositol and phosphate into the blood or urine.

Subsequently, investigators who assumed that phytate is absorbed and can be detected in the blood or urine (Grases et al., 2000) were unable to detect phytate in serum or urine despite using a highly specific and sensitive mass spectrometry (Letcher et al., 2008). These studies support the conclusion that phosphate or *<u>myo</u>*-inositol generated by the hydrolysis of phytate hydrolysis is absorbed rather than phytate, which is unabsorbable. Numerous explanations account for the impermeability of intestinal membranes to phytate (Reddy, 2002). Phytate possesses six phosphate groups with strong negative charges at physiological pH, which suggests that phytate cannot cross the lipid bilayer of the plasma membrane without a specific receptor (Ozaki et al., 2000). In support of this possibility, we found that phytate is only engulfed by hepatic Kupffer cells and tissue macrophages *in vivo* when we intravenously injected iron-containing phytate particles into live mice (Kim et al., 2017; Lee et al., 2018), indicating that phytate cannot penetrate membranes of epithelial cells in the intestine. Together, these findings suggest that phytate is not absorbed in the intestine and that only products generated through the hydrolysis of phytate, phosphate, or *myo*-inositol can be absorbed in the intestine of rats fed phytate-containing low-Ca^2+^ diets.

Here we demonstrate that phytate can act as a dual demon in healthy normal rats by adversely affecting Ca^2+^ and phosphate homeostasis through inhibition of intestinal Ca^2+^ absorption while increasing intestinal phosphate overload, depending on the Ca^2+^ content of diets. This may explain why rats fed the HP-LCa^2+^ diet exhibited high levels of PTH and 1,25(OH)_2_D_3_, as well as low levels of FGF23 and αKlotho. Moreover, our findings are consistent with data from a large population-based study of subjects with preserved renal function without CKD, which showed that PTH levels increase earlier than FGF23 levels as kidney function decreases and that high FGF23 levels are more closely associated with low levels of 1,25(OH)_2_D_3_ than with excessive renal phosphate wasting (Dhayat et al., 2016). Thus, our results suggest that high levels of 1,25(OH)_2_D_3_ may suppress the expression of calvarial FGF23 in rats fed the HP-LCa^2+^ diet, in accordance with an earlier study showing that Ca^2+^ deficiency is highly associated with low levels of FGF23 (Rodriguez-Ortiz et al., 2012). Moreover, Ca^2+^ deficiency increases the levels of 1,25(OH)_2_D_3_, which leads to low levels of FGF23 in Ca^2+^ deficiency states (Clements et al., 1987).

However, previous studies that investigated the effects of phytate on kidney stones in rats report conflicting results, which we ascribe to differences in dietary compositions, particularly to the ratio of Ca^2+^/phytate and analytical laboratory methods (Grases et al., 2000). Specifically, the AIN-76A purified rodent diet containing 1% phytate prevents the development of kidney stones compared with the AIN-76A controls (Grases et al., 2000). Further, Grases et al. (17) concluded that rats fed the AIN-76A diet developed mineral deposits at the corticomedulary junction from Alizarin Red S staining, whereas Lien et al. (Lien et al., 2001) did not find an increased incidence or severity of renal tubular mineralization in rats fed the AIN-76A or AIN-93G diet. Consistent with this, we found that rats fed the AIN-93G control diet did not have nephrocalcinosis. Moreover, we found that an AIN-93G-based HP-LCa^2+^ diet induced biochemical and urinary abnormalities associated with kidney disease as well as histopathological defects in the kidney. Phytate in the AIN-96-G diet caused a dramatic time- and concentration-dependent increase in PTH levels, leading to PTH-induced impairment of the renal reabsorption of Ca^2+^ and phosphate and predisposed rats to the development of hypophosphatemic rickets and nephrocalcinosis.

In conclusion, we demonstrate here that a high-phytate low-Ca^2+^ diet is a risk factor for nephrocalcinosis, renal inflammation, and fibrosis as well as renal phosphate wasting disorders and bone loss in rats. Our studies provide an animal model for studying the pathogenesis of crystal nephropathies and bone loss associated with renal phosphate wasting and further suggest that the restriction of phytate intake may reduce the risks of developing crystal nephropathies.

## Materials and Methods

### Care and feeding of rats

Animal studies were approved by the Center for Animal Care and Use and were performed according to institutional ethics and safety guidelines (LCDI-2008-0019, LCDI-2010-0068, and LCDI-2018-0060). Three-week-old male (60–70 g) and female (50–55 g) Sprague–Dawley rats (Orient Bio, Seoul, Korea) were housed three per cage in a 12:12-h light-dark cycle with an ambient temperature maintained at 22°C ± 2°C. The rats were fed AIN-93G (Dyets Inc., Bethlehem, PA, USA) for 1 week and were randomly assigned to control AIN-93G (0% phytate), AIN-93G with 1%, 3%, or 5% sodium phytate (P-8810, Sigma-Aldrich, St. Louis, MO, USA) or AIN-93G with 3% sodium phytate with an additional 0.5% Ca^2+^ (n = 6–12 per group) diet for 12 weeks. All diets had equivalent percentages of carbohydrate, protein, fat, and minerals. Pathogen-free water and food were available *ad libitum*. Food intake and body weight were recorded weekly.

### Biochemical assays

The rats were fasted overnight every 2–3 weeks and anesthetized with isoflurane. Tail vein blood samples were stored at 4°C for 10–20 min and serum was frozen at −80°C. At the end of the experiment, the rats were fasted for 14–16 h before receiving isoflurane. Abdominal aorta blood was obtained, and urine was collected over 24 h. The kidneys and femora were dissected and stored at −80°C. Serum biochemical analysis was performed (Model AU-480; Beckman Coulter, Fullerton, CA, USA). Rat intact PTH (Immutopics, San Clemente, CA, USA), human intact PTH (Abcam), soluble RANKL (Immundiagnostik, Bensheim, Germany), osteoprotegerin (Alpco Immunoassay, Salem, NH, USA), rat intact and C-terminal FGF23 (Elabscience, Houston, TX, USA), and human intact FGF23 (Elabscience) were measured using enzyme-linked immunosorbent assay. Serum 25(OH)D_3_ concentrations were determined by chemiluminescence microparticle immunoassay (Abbott, Abbott Park, IL, USA) and serum 1,25(OH)_2_D_3_ concentrations were determined by radioimmunoassay (DIAsource, Belgium) in the biochemistry laboratory at Green Cross Labcell (Yongin, Korea). Creatinine clearance (Cl) was calculated as Cl (ml/min) = (urine [creatinine] × urine volume × body weight)/(serum [creatinine] × 24 × 60). The fractional excretion of Ca^2+^ or phosphate was calculated as the %FE_Ca or_ Pi = urine [Ca^2+^ or phosphate] × serum [creatinine])/(urine [creatinine] × serum [Ca^2+^ or phosphate]) × 100. TmP/GFR and TmCa^2+^/GFR were calculated in accordance with the Walton and Bijvoet nomogram (Walton and Bijvoet, 1975) using a 24-h urine sample and a serum sample obtained 16 h after urine collection and fasting.

### Fecal mineral content analysis

Fecal samples were collected from rats after 2 or 4 weeks of either the control or phytate-supplemented diet. Rats were placed in grid cages over clean trays to collect samples. Food was removed from the cages during collection to avoid contamination. All samples were stored at −80°C. Ca^2+^, magnesium, iron, and phosphate were quantified on an inductively coupled plasma optical emission spectrometer (Optima 5300 DV; PerkinElmer, Waltham, MA, USA),. Fecal samples were dried at 60°C for 48 h, and 1.0 g was heated at 560°C for 4 h. After cooling to room temperature, 1 mL of 35.5% HNO_3_ was added and samples were heated at 560°C for an additional 2 h. Samples were then exposed to 4 mL 17.5% HCl at room temperature for 2 h before dilution in ultra-pure water. Fecal phytate was sequentially extracted with a HCl-HF acid solution, followed by an alkaline extraction (NaOH and EDTA), as previously described (Ray et al., 2012). The phytate content of fecal extracts was quantified by high-performance ion chromatography (ICS-3000; Dionex).

### Kidney histopathology

Hematoxylin and eosin (HE)-stained kidney sections (n = 64) from eight male and eight female rats were blindly examined by one author (CJB). Ca^2+^ was stained with Alizarin Red S and von Kossa. Sections obtained from females were stained with Periodic acid–Schiff. Parameters were scored semiquantitatively based on previous work (Lalioti et al., 2006). The measured parameters included the presence and severity of mineralization (0 = negative or within normal limits, 1 = minimal, 2 = mild, 3 = moderate, 4 = marked, and 5 = severe), inflammation, tubular casts, tubular ectasia, interstitial fibrosis, tubular degeneration/regeneration, and glomerular hypertrophy. Individual scores were added together for each kidney, with a maximum score of 40.

### In vivo magnetic resonance imaging (MRI)

We used a 9.4 T animal MRI scanner (Biospec 94/20 USR; Bruker Biospin, Ettlingen, Germany) with a transmit-only volume coil for excitation and phased-array four-channel surface coil for signal reception (Bruker Biospin). Animals were anesthetized using isoflurane. MRI scans were performed using the following respiratory-gated, fat-suppressed gradient echo sequence: TR/TE, 5000/28.5 ms; FA, 180°; FOV, 70 × 60 mm^2^; matrix size, 384 × 384; TH, 1.0 mm; 27 slices (no gap); and NEX, 4. Axial images were acquired for anatomic reproducibility and histopathological correlation.

### Bone mineral density (BMD)

The right femora samples were harvested from eight mice per group after 12 weeks on either control or phytate-supplemented diet. *Ex vivo* BMD, bone mineral content, and femoral lengths were measured by peripheral dual-energy X-ray absorptiometry (pDEXA) (Sabre™; Norland Medical Systems Corp., Fort Atkinson, WI, USA) with pDEXA Sabre software (version 3.9.4; Norland). Scans of femora (4 × 4 cm^2^) were performed at 3 mm/s and resolution of 0.2 × 0.2 mm. BMD was computed using dual X-ray beam attenuation (28 and 48 KeV), and BMD histogram averaging widths were 0.02 g/cm^2^. *In vitro* precision, expressed as the coefficient of variation (CV = 100 × standard deviation/mean), was calculated by the BMD of a phantom with a nominal density of 0.929 g/cm^2^.

### μ-computed tomography (µ-CT) analysis

The femora were isolated from soft tissue, fixed overnight in 10% formalin, and analyzed by high-resolution μ-CT (SkyScan1172 *in vivo* CT; Bruker). Imaging (NRecon ver. 1.6; Bruker), data analysis (CTAn ver. 1.11; Bruker), and three-dimensional visualization (μ-CTVol ver. 2.2) were performed on 500 slices per scan at 70 kVp, 141 μA, and 11.55 μm pixel^−1^. Cross-sectional images were obtained for three-dimensional histomorphometry. A 1.2-mm sample of metaphyseal secondary spongiosa was imaged. Cross-sectional images (500 images per specimen) of distal femora were used for two-dimensional and three-dimensional morphometry.

### Bone histomorphometry

The right femora were fixed in 10% formalin for 48 h, decalcified with 8% hydrochloric acid/formic acid for 2 weeks, and embedded in paraffin. Longitudinally oriented 2.5-μm thick bone sections (including metaphyses and diaphyses) were processed for HE, Toluidine Blue O, and Masson’s trichrome stains. The left femora were fixed in formalin for 48 h, decalcified in 10% EDTA (pH 7.0) for 4–5 weeks at 4°C, and embedded in paraffin for tartrate-resistant acid phosphatase staining. Sections were viewed using a Zeiss Axio Imager Z1 microscope (Carl Zeiss, Oberkochen, Germany). Histomorphometric measurements were made at the center of the metaphyses. Images were captured with 3DHISTECH (Budapest, Hungary). Bone sections (2.5-μm thick) were stained with Masson’s trichrome for osteoids.

### Rat bone marrow osteoclasts

Bone marrow cells were isolated using a modified method (Kobayashi et al., 2000). Tibiae and femora were isolated from rats after 12 weeks on either control or phytate-supplemented diet. Bone ends were removed and the marrow was flushed by slowly injecting serum-free alpha minimal essential medium (αMEM; Invitrogen, Carlsbad, CA, USA) through a sterile 25-gauge needle into one end of the bone. Marrow cells were collected into tubes, washed twice with αMEM, and cultured in αMEM with 10% heat-inactivated fetal bovine serum (Invitrogen). Cells were cultured in 24-well plates (1 × 10^5^ /well) with 30 ng/mL of macrophage colony-stimulating factor (M-CSF) for 72 h. Differentiation of M-CSF-dependent bone marrow osteoclast precursors was induced with 30 ng/mL receptor activator of nuclear factor kappa-B ligand (RANKL) (R&D Systems, Minneapolis, MN, USA) and 30 ng/mL M-CSF for 4 days. Differentiated osteoclasts with >3 nuclei were identified as tartrate-resistant acid phosphatase-positive (Leukocyte Acid Phosphatase Assay, Sigma).

### Western blotting and immunohistochemistry (IHC)

The primary antibodies used for Western blotting were polyclonal CYP27B1 (M-100, 1:200; Santa Cruz Biotechnology, Inc. Santa Cruz, CA, USA), CYP24A1 (S-20, 1:200; Santa Cruz), and VDR (C-20, 1:200; Santa Cruz), and monoclonal β-actin (AC-15, 1:1000; Sigma), FGF23 (bs-5768R, 1:200; Bioss), αKlotho (NBP1-76511, 1:500; Novus Biologicals), NHERF1 (A-7, 1:200; Santa Cruz), and SLC34A1 (ab182099, 1:500; Abcam). Antigen retrieval for IHC was performed in Tris/EDTA (pH 9.0) for 2–3 min. IHC was performed with the following polyclonal antibodies (Santa Cruz) at 25°C for 2 h: CYP27B1 (M-100, 1:50), CYP24A1 (S 20, 1:50), VDR (D-6, 1:50), RANKL (N-19, 1:50), or osteoprotegerin (N-20, 1:50), as described previously (Kang et al., 2017). Horseradish peroxidase-conjugated secondary antibodies were detected with 3,3,-diamino-benzidine tetrahydrochloride substrate (Dako, Glostrup, Denmark). All sections were counterstained with hematoxylin (Dako).

### qRT-PCR analyses

Total RNA was extracted from whole kidney lysates with Trizol (Invitrogen). cDNA synthesis and PCR amplification were performed using PrimeScript First-Strand cDNA Synthesis kit (TaKaRa Bio, Tokyo, Japan) with 1 μg total RNA plus random hexamers. Real-time RT-PCR was performed using SYBR green (Takara) in BioRad CFX384. All expression values were normalized to the *cyclophilin A* mRNA level. Primer sequences are listed in **table supplement 2**.

### Phytate pilot study in humans

A noncomparative pilot study was approved by the Institutional Review Board of Dongguk University Ilsan Hospital (IRB No. 2014-117). All procedures were performed in accordance with the Declaration of Helsinki. The study was registered with Clinical Research Information Service (registration number KCT0003144) in accordance with the WHO International Clinical Trials Registry Platform. This study aimed to evaluate the effects of phytate intake on mineral metabolism in humans between January and February 2015. All participants provided written informed consent. In total, six premenopausal women aged between 23 and 34 years with a body mass index between 17.7 and 25.8 kg/m^2^ were recruited. The exclusion criteria were as follows: bone and mineral metabolism disorders (e.g., hyperparathyroidism, hypoparathyroidism, osteomalacia), previous surgical history which could affect bone and mineral metabolism (e.g., parathyroidectomy, thyroidectomy, gastrectomy), use of any medication that could affect bone and mineral metabolism (e.g., estrogen, bisphosphonates, selective estrogen receptor modulators, calcitonin), use of Ca^2+^ and vitamin D supplementation within the past 3 months, use of corticosteroids within the past 1 month, or acute or chronic renal failure; exhibited abnormal liver function; or were pregnant or lactating. All participants completed a 12-day trial consisting of three successive diet cycles containing (1) polished rice (low phytate content, 0.35%); (2) brown rice (medium phytate content, 1.07%); and (3) brown rice mixed with rice bran (high phytate content, 2.17%). Each diet cycle lasted for 4 days. During each day of the study, all participants were provided with breakfast, lunch, dinner, and a snack. Every meal was packed in a lunch box and delivered each day, and participants were not allowed to consume any other food during the trial except for the food provided by study investigators. All meals were identical across the three diet cycles except for the phytate content. Water consumption was allowed *ad libitum*. No medications or supplements that affected mineral metabolism were allowed during the course of the study. A blood sample was collected from each participant at the beginning of the trial and at the end of each diet cycle. Urine samples of 24 h were collected before the trial and during the fourth day of each diet cycle.

### Statistical analysis

Sample sizes for all experiments were based on preliminary results and previous experience of conducting related experiments. Power calculations were not used to determine sample sizes. Animals for each group of experiments were randomly assigned. No animals were excluded from statistical analysis, and investigators were blinded to the study condition when scoring histopathological samples. Unless noted otherwise, graphical presentations show the mean ± standard deviation (SD). Comparisons across groups were conducted using either unpaired Student’s *t* test or one- or two-way ANOVA with Dunnett’s or Tukey’s post hoc multiple comparison testing. Statistical analyses were performed using GraphPad Prism 8.2 (GraphPad Software, Inc., San Diego, CA).

## Acknowledgments

The authors thank Dr. S. Dhe-Paganon and S. Shoelson at Harvard Medical School for their valuable suggestions regarding the manuscript.

## Supplemental figures

**Figure supplement 1.**
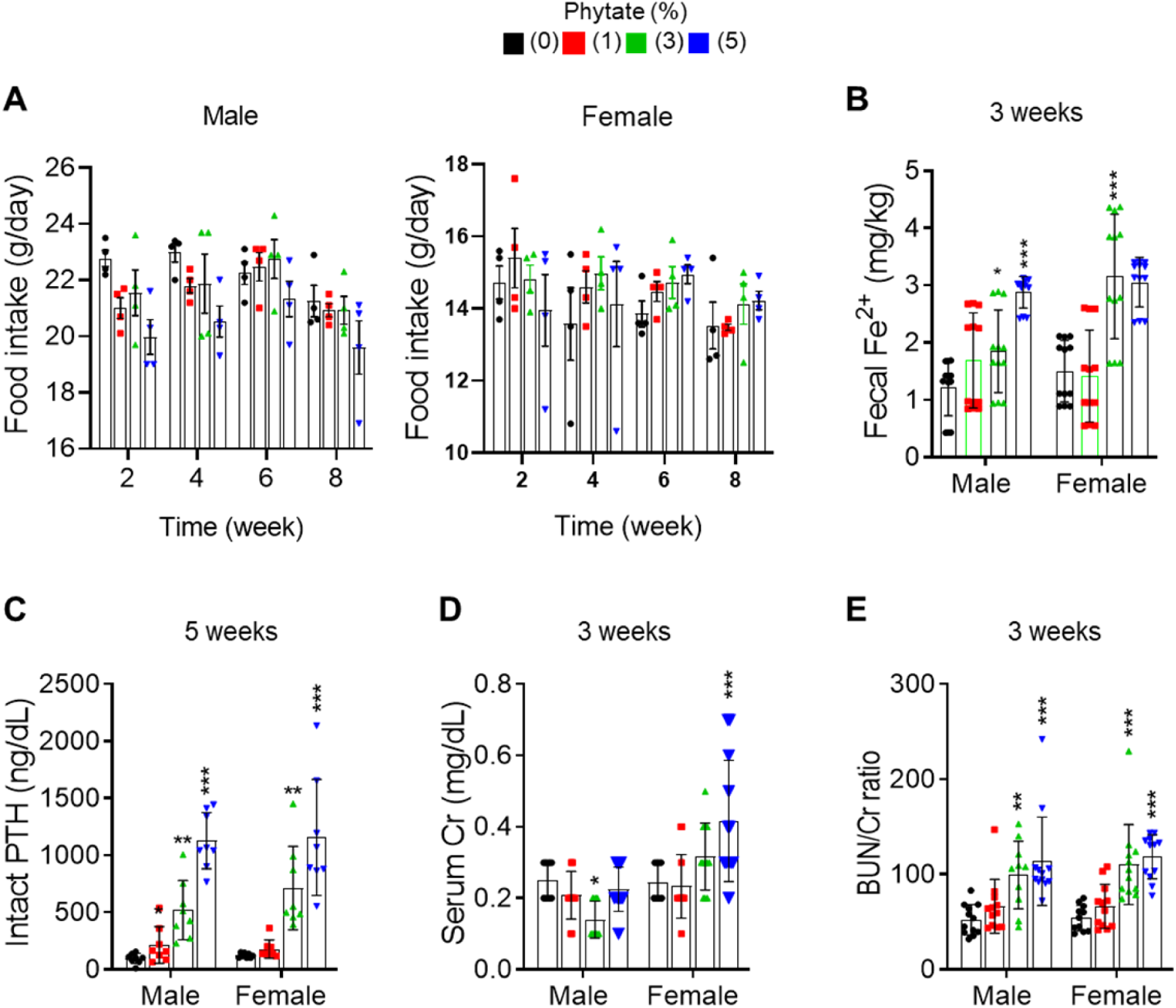
Metabolic characteristics of Sprague-Dawley rats fed control and phytate-supplemented diets. (**A)** Food intake of rats fed control or phytate-supplemented diets. (**B**) Fecal excretion of Fe^3+^ in rats fed control or phytate-supplemented diets. (**C-E)** The serum levels of intact PTH (**C**), creatinine (**D**), and BUN/Cr ratio (**E**) in rats after 3, or 5 weeks on a control or phytate-supplemented diet. All comparisons were analyzed by one-way ANOVA, *P* values were corrected using Dunnett’s multiple comparisons test. All data are presented as the mean ± SD for each group (n = 8–12 per group). *P < 0.05 compared with controls.

**Figure supplement 2.**
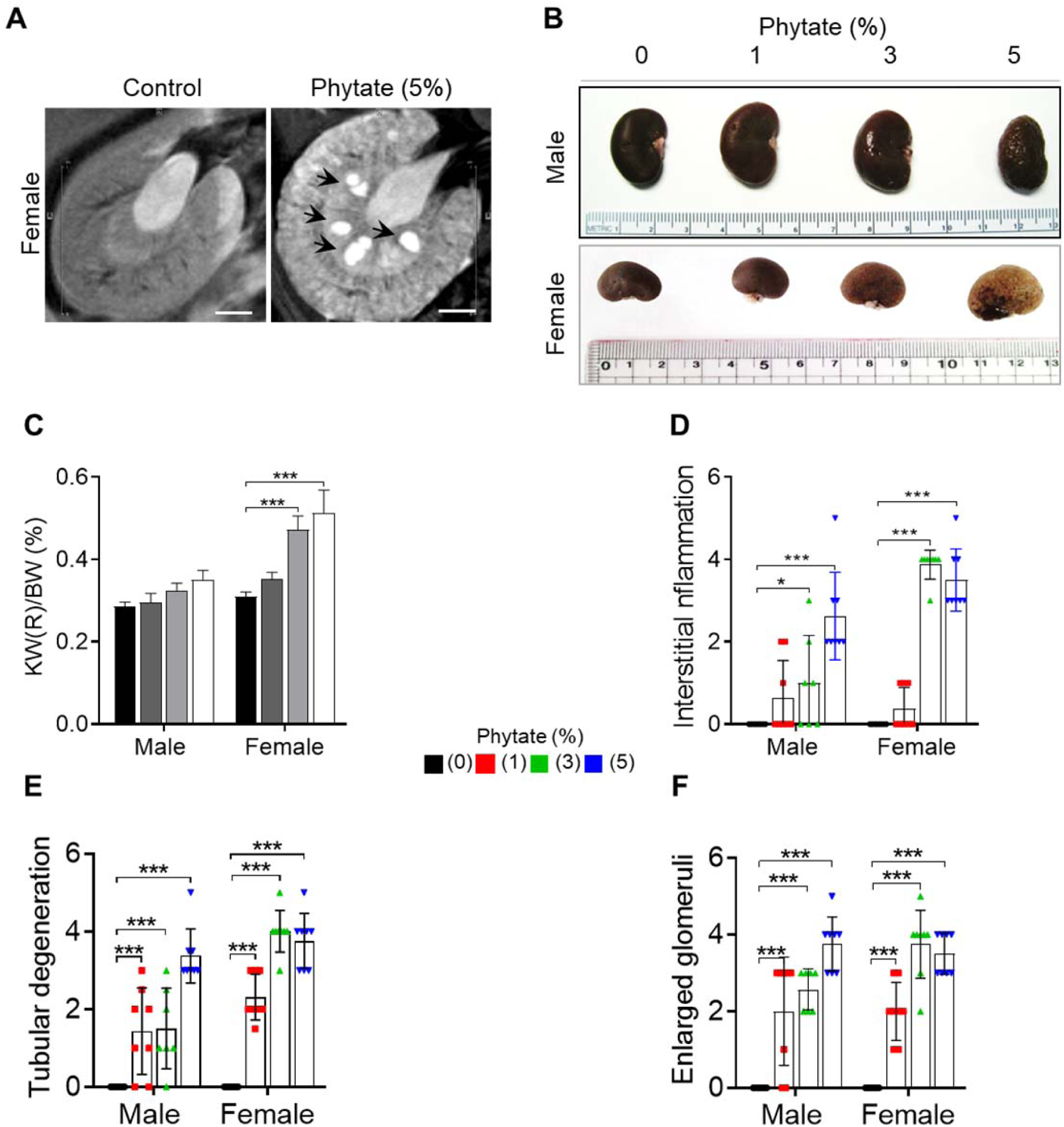
Histopathological features of the kidneys of phytate-fed rats. (**a)** Representative T2-weighted magnetic resonance images (MRIs) of the kidneys of female rats fed either a control or 5% phytate diet. (**b-c)** Representative gross morphological features of the kidneys (**b**) and kidney weight (%) relative to body weight (**c**, KW/BW) of rats fed control and phytate-supplemented diets. (**d-f)** The renal histopathological scores for interstitial inflammation (**d**), tubular degeneration (**e**), and enlarged glomeruli (**f**) from rats fed control and phytate-supplemented diets. All comparisons were analyzed by one-way ANOVA, *P* values were corrected using Dunnett’s multiple comparisons test. All data are presented as the mean ± SD for each group (n = 8). *P < 0.05 compared with controls.

**Figure supplement 3.**
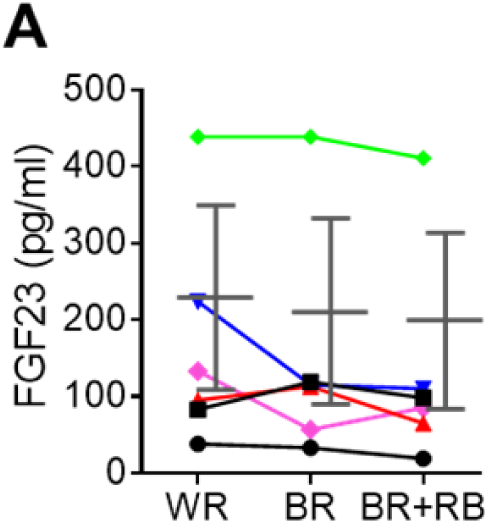
High-phytate diets dysregulate mineral and phosphate metabolism in humans. (**A)** Time course analysis of serum levels of intact FGF23 for three consecutive phytate doses. Statistical significance were tested by repeated measure ANOVA between subjects and groups. Data are presented as the mean ± SD for each dose (n= 6 per dose).

**Figure supplement 4.**
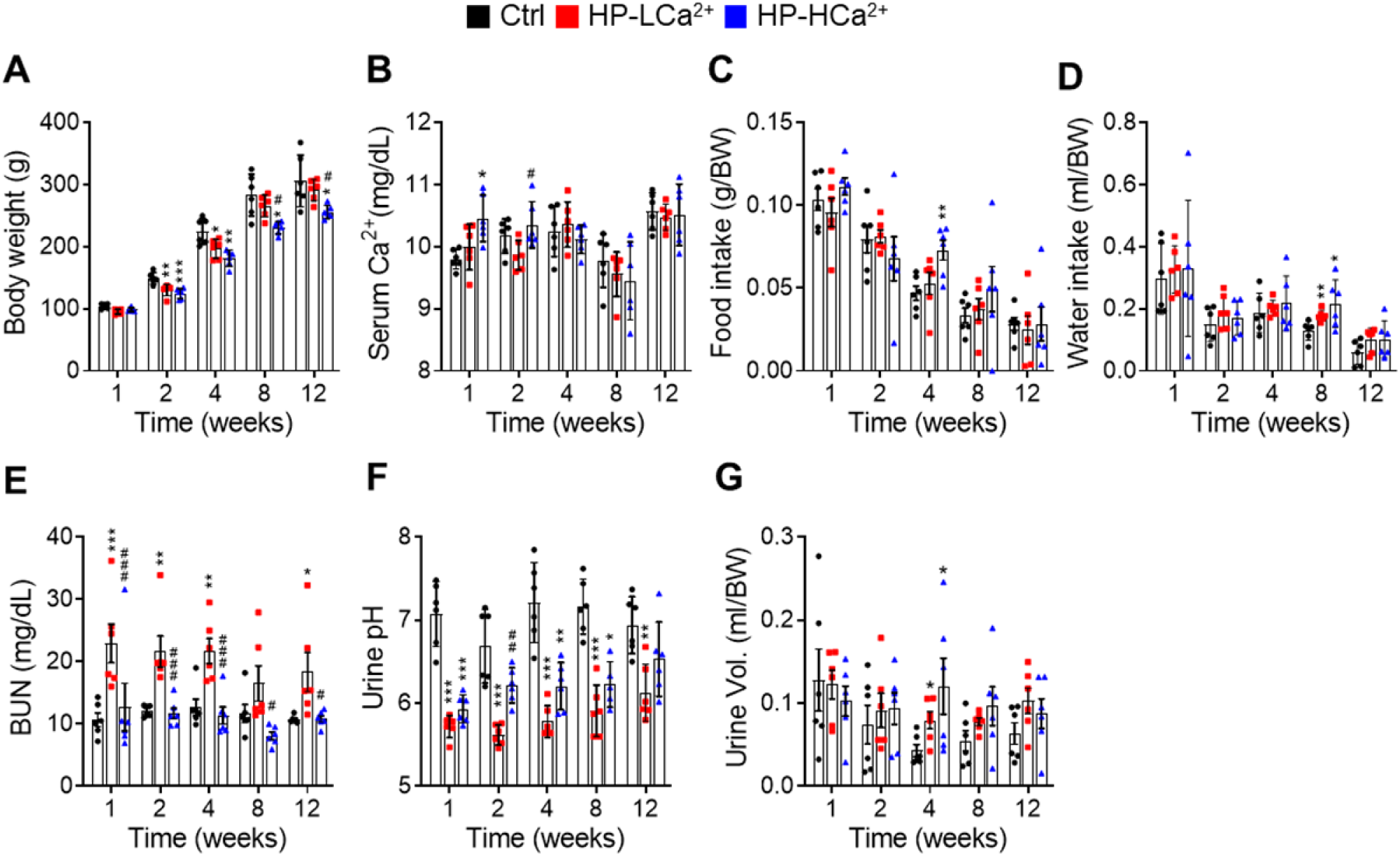
Ca^2+^ supplementation protects rats fed phytate from crystal nephropathies. **(A-D)** Time course analysis of kidney weight (%) relative to body weight (KW/BW, **A**), representative gross morphological features of kidneys (**B**,**C**) and Alizarin red S staining (**D-F**) of kidney sections from rats fed control, HP-LCa^2+^, and HP-LCa^2+^ diets. (**G)** The nephrocalcinosis area was quantified from the Alizarin Red S staining of kidney sections using ImageJ software (NIH, Bethesda, MD, USA). The data are presented as the mean ± standard error of the mean (SEM) for each group (n =6 per group).

**Figure supplement 5.**
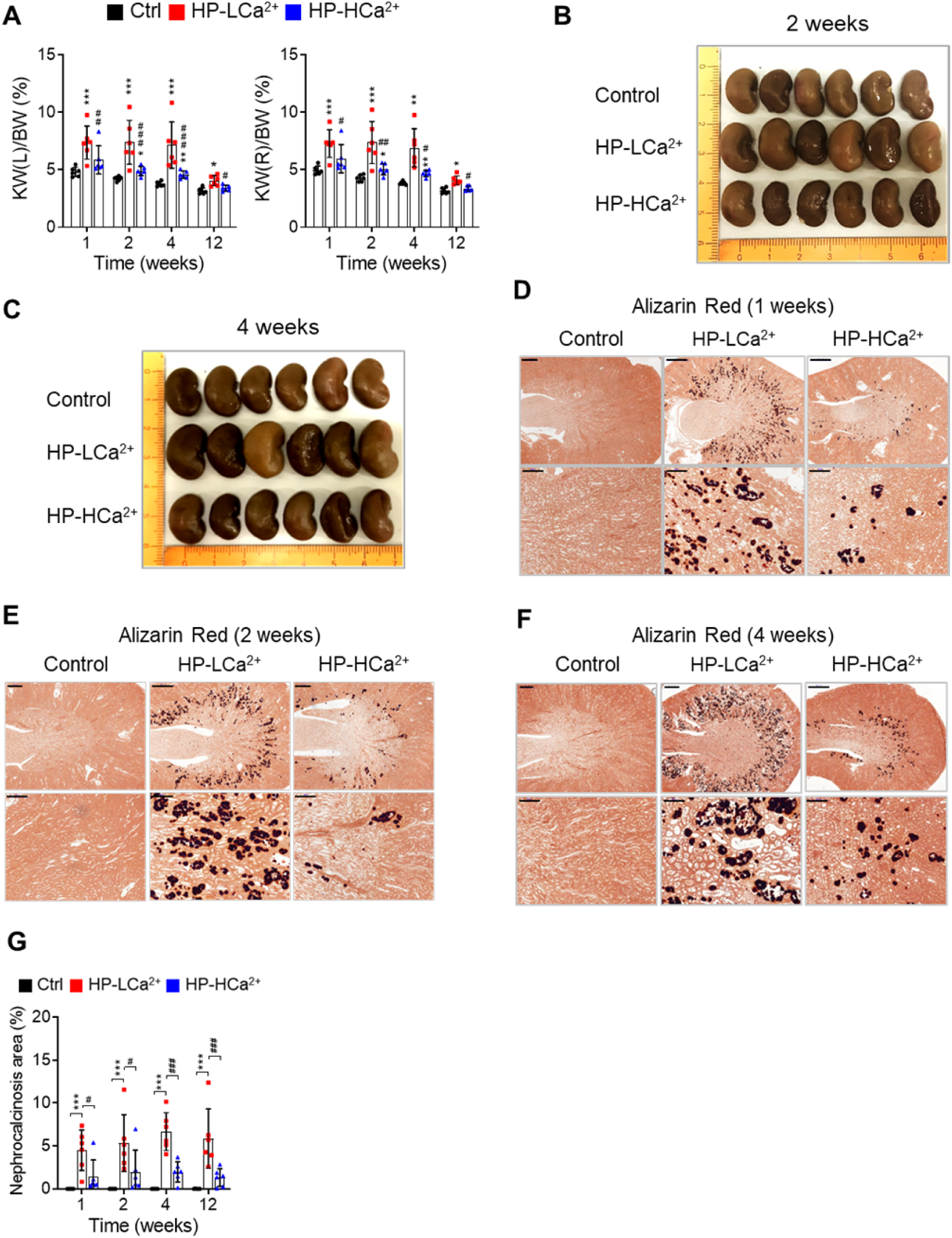
Ca^2+^ supplementation protects rats fed phytate from crystal nephropathies. **(a-d)** Time course analysis of kidney weight (%) relative to body weight (KW/BW, **a**), representative gross morphological features of kidneys (**b**,**c**) and Alizarin red S staining (**d-f**) of kidney sections from rats fed control, HP-LCa^2+^, and HP-LCa^2+^ diets. (**g)** The nephrocalcinosis area was quantified from the Alizarin Red S staining of kidney sections using ImageJ software (NIH, Bethesda, MD, USA). All comparisons were conducted using two-way ANOVA with Tukey’s post hoc multiple comparison testing. All data are presented as the mean ± SD of each group (n=6 per group). *P < 0.05, **P < 0.01, ***P < 10^−3^, compared with controls. ^#^P<0.05, ^##^P<0.01, ^###^P<10^−3^, compared with a HP-LCa^2+^ diet groups.

**Figure supplement 6.**
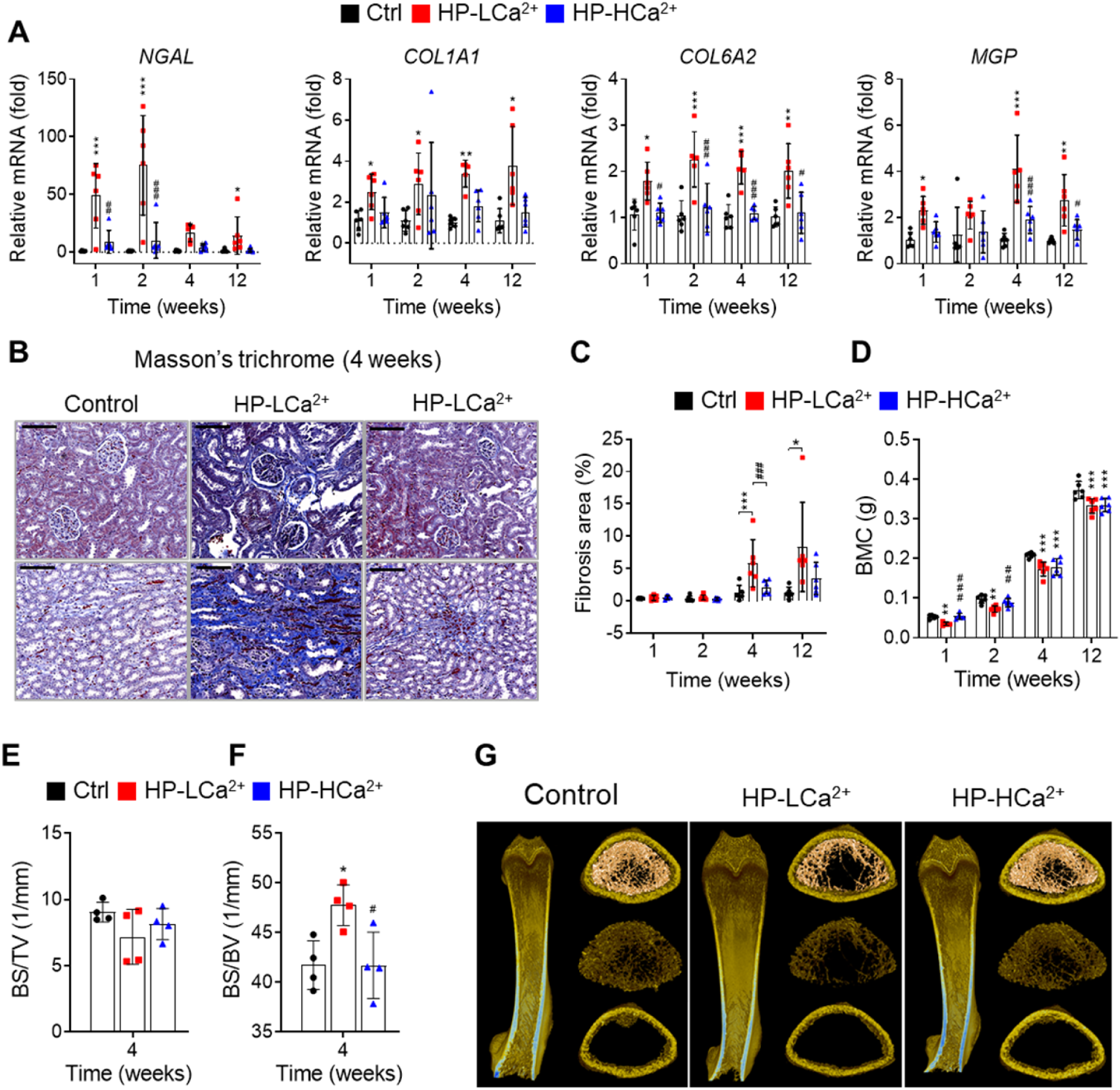
Ca^2+^ supplementation protects phytate-fed rats from renal fibrosis and skeletal abnormalities. (**A)** Time course analysis of renal expression of *NGAL, COL1A1, COL6A2*, and *MGP* in rats fed control, HP-LCa^2+^, and HP-HCa^2+^ diets for 12 weeks. (**B)** Masson’s trichrome staining from rats fed control, HP-LCa^2+^, and HP-HCa^2+^ diets. (**C)** Renal fibrosis area was quantified from Masson’s trichrome staining of kidney sections using ImageJ software. (**D-H)** Time course analysis of BMC (**D**), bone surface density (**E**, BS/TV), and specific bone surface (**F**, BS/BV) from rats fed control, HP-LCa^2+^, and HP-HCa^2+^ diets. Representative three-dimensional reconstructed μ-CT images of distal femora (**G**) from rats fed control, HP-LCa^2+^, and HP-HCa^2+^ diets. All comparisons were conducted using two-way ANOVA with Tukey’s post hoc multiple comparison testing. All data are presented as the mean ± SD of each group (n =4-6 per group). *P < 0.05, **P < 0.01, ***P < 10^−3^, compared with controls. ^#^P<0.05, ^##^P<0.01, ^###^P<10^−3^, compared with a HP-LCa^2+^ diet groups.

**Figure supplement 7.**
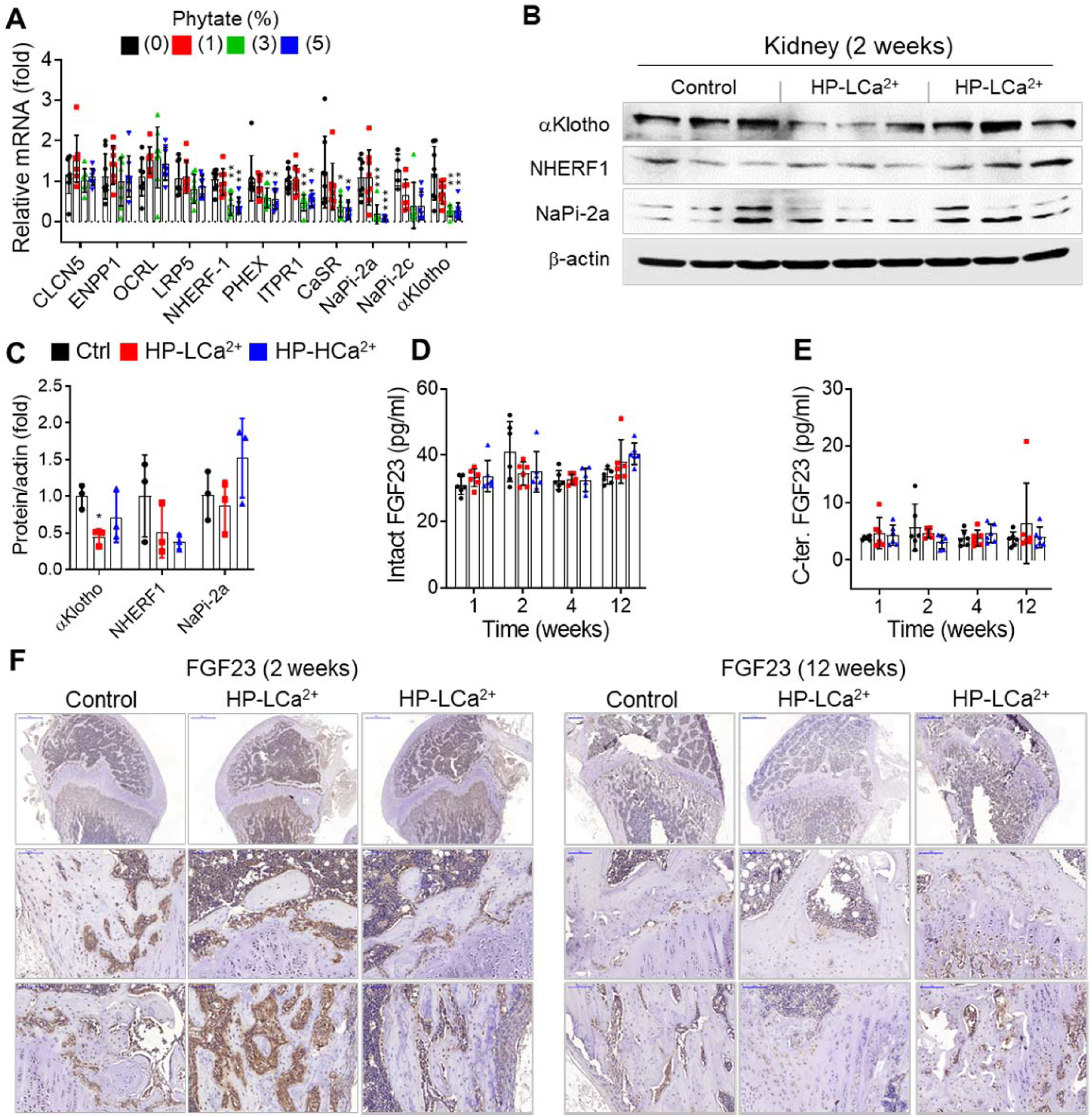
High phytate-induced renal phosphate wasting is independent of FGF23. (**A)** The renal expression of down-regulated genes associated with Ca^2+^ and phosphate homeostasis in rats fed control or phytate-supplemented diets (n = 8 per group). (**B)** Results of immunoblot analyses and protein levels of renal aKlotho, NHERF1, and NaPi-2a after 2 weeks on rats fed control, HP-LCa^2+^, and HP-HCa^2+^ diets. (**C)** The levels of renal proteins were quantified using ImageJ software (n=6 per group). (**D-E)** Time course analysis of serum levels of intact FGF23 (**D**) and C-terminal FGF23 (**E**) in rats fed control, HP-LCa^2+^, and HP-HCa^2+^ diets. (**F)** Representative immunohistochemical staining of EDTA-decalcified femur sections for FGF23 from rats fed control, HP-LCa^2+^, and HP-HCa^2+^ diets. All data are presented as the mean ± SD of each group (n =3-8 per group). *P < 0.05, **P < 0.01, ***P < 10^−3^, compared with controls.

**Table 1.**
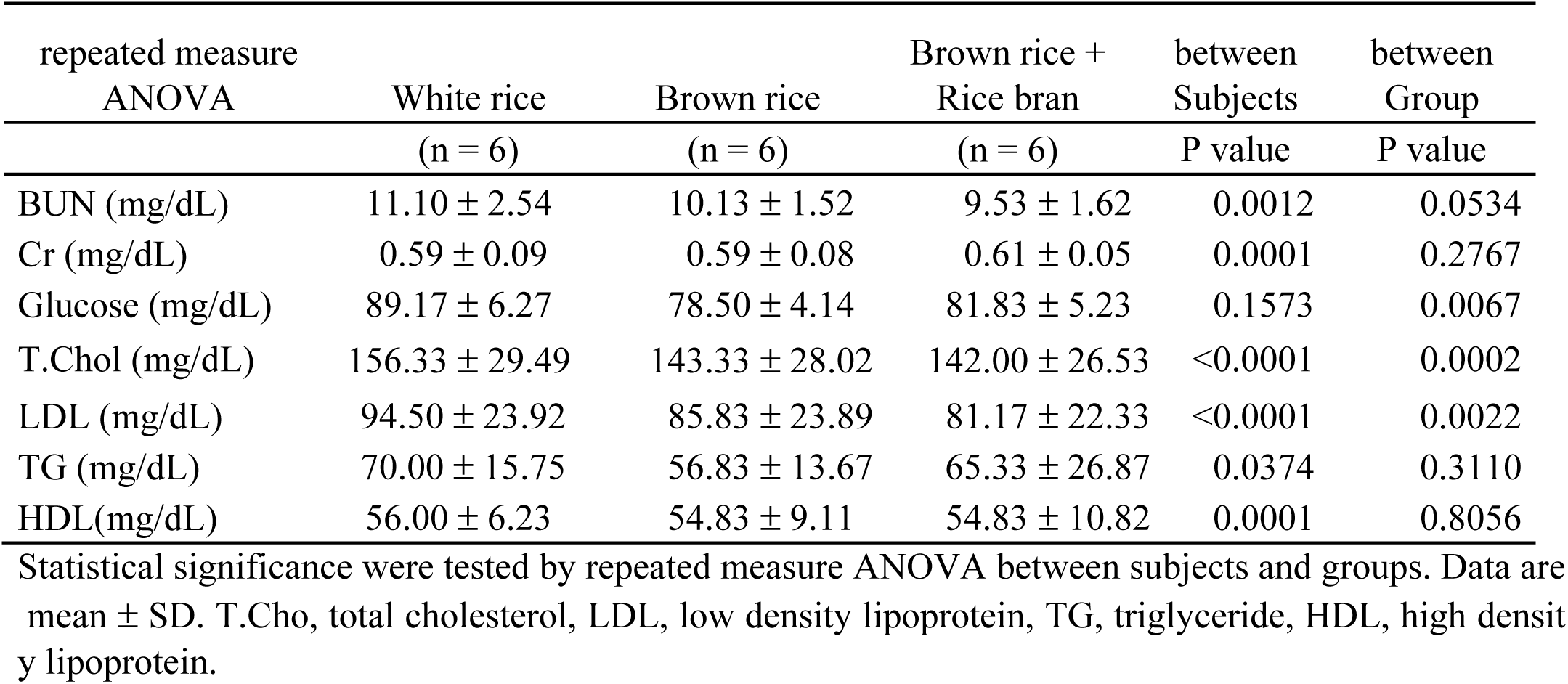
Biochemical properties during three consecutive diets.

| repeated measure<br>ANOVA | White rice | Brown rice | Brown rice +<br>Rice bran | between<br>Subjects | between<br>Group |
| --- | --- | --- | --- | --- | --- |
|  | (n = 6) | (n = 6) | (n = 6) | P value | P value |
| BUN (mg/dL) | 11.10 ± 2.54 | 10.13 ± 1.52 | 9.53 ± 1.62 | 0.0012 | 0.0534 |
| Cr (mg/dL) | 0.59 ± 0.09 | 0.59 ± 0.08 | 0.61 ± 0.05 | 0.0001 | 0.2767 |
| Glucose (mg/dL) | 89.17 ± 6.27 | 78.50 ± 4.14 | 81.83 ± 5.23 | 0.1573 | 0.0067 |
| T.Chol (mg/dL) | 156.33 ± 29.49 | 143.33 ± 28.02 | 142.00 ± 26.53 | <0.0001 | 0.0002 |
| LDL (mg/dL) | 94.50 ± 23.92 | 85.83 ± 23.89 | 81.17 ± 22.33 | <0.0001 | 0.0022 |
| TG (mg/dL) | 70.00 ± 15.75 | 56.83 ± 13.67 | 65.33 ± 26.87 | 0.0374 | 0.3110 |
| HDL(mg/dL) | 56.00 ± 6.23 | 54.83 ± 9.11 | 54.83 ± 10.82 | 0.0001 | 0.8056 |

Statistical significance were tested by repeated measure ANOVA between subjects and groups. Data are mean ± SD. T.Cho, total cholesterol, LDL, low density lipoprotein, TG, triglyceride, HDL, high densit y lipoprotein.

**table Supplement 2.**
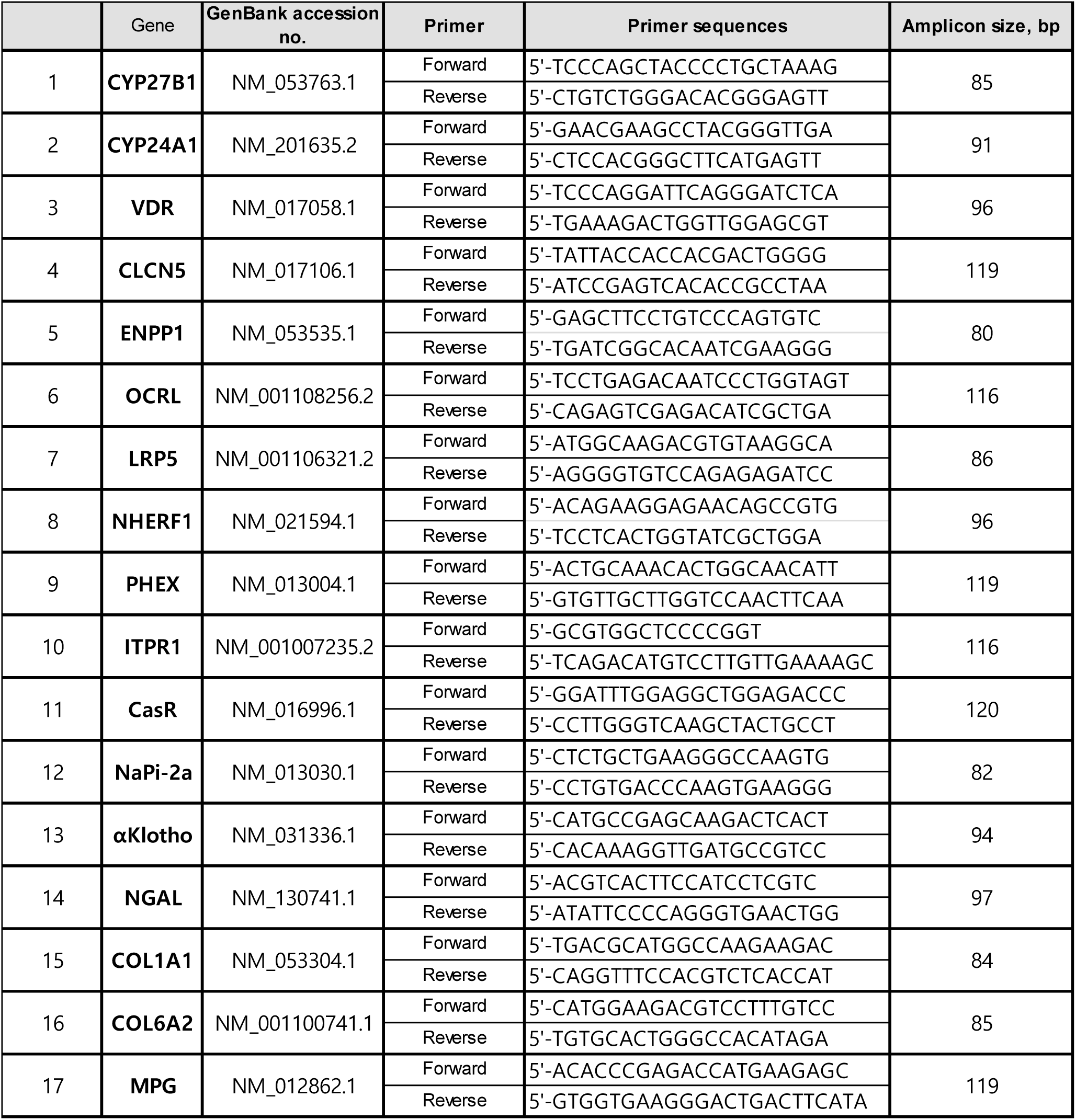
The qRT-PCR primer lists.

